# A Cryptic Interfacial Pocket Uncovered in Full CRL4^CRBN^–IKZF3 Ubiquitylation Complex Enhances IMiD Efficacy

**DOI:** 10.1101/2025.06.08.658527

**Authors:** Zhiheng Deng, Huasong Ai, Qiang Shi, Jiawei Liang, Zaozhen He, Jiqing Zheng, Haoxiang Li, Liying Zhang, Wei He, Shixian Tao, Qingyun Zheng, Wei He, Man Pan, Lei Liu

## Abstract

Immunomodulatory imide drugs (IMiDs) redirect the CUL4–RBX1–DDB1–CRBN (CRL4^CRBN^) ligase to ubiquitylate and degrade disease-linked proteins, but the full picture of how IMiDs glue the neosubstrate within the whole CRL4^CRBN^ complex remains unknown. We determined cryo-electron microscopy structures of eight approved or clinical IMiDs in full CRL4^CRBN^ ubiquitylation complexes at resolutions up to 3.4 Å, and revealed how these structurally distinct IMiDs exploit the structural plasticity of CRL4^CRBN^ to organize conformationally conserved and compact active ubiquitylation assemblies with the neosubstrate IKZF3. Four “next-generation” IMiDs were found to additionally engage a cryptic gluing-driven interfacial (GDI) pocket at the non-degron zinc finger 3 (ZF3) of IKZF3, which contributes to their enhanced efficacy and neosubstrate specificity. The identification of cryptic gluing-driven pockets formed only in full ubiquitylation complexes provides new structure-directed design opportunities for IMiDs to improve therapeutic efficacy.

## Introduction

Transcending its tragic clinical origins, the teratogenic drug thalidomide ^1^was discovered to be repurposable for treating diseases with historically limited therapeutic options, particularly in oncology and hematology ^2^. This fortuitous finding has both paralleled and driven myriad innovations in the history of pharmaceuticals. Thalidomide-inspired immunomodulatory imide drugs (IMiDs), including lenalidomide and pomalidomide, function through cereblon (CRBN) as part of the CRL4^CRBN^ E3 ligase complex ^3-5^ and have demonstrated remarkable clinical efficacy in treating multiple myeloma ^6^. IMiDs recruit the CRL4^CRBN^ E3 ligase complex to ubiquitylate and facilitate the subsequent degradation of specific cellular substrates, including the Ikaros family transcription factors, IKZF1 and IKZF3 ^7-9^, and casein kinase 1α (CK1α) ^10^, leading to tumor suppression and immune modulation. These agents demonstrate high efficacy, with lenalidomide generating over $12 billion in annual sales and pomalidomide showing improved survival in relapsed/refractory patients ^6,11^.

Decades of studies have elucidated how IMiDs rewire protein–protein interactions (PPIs) to achieve therapeutic consequences ^12-15^. A plethora of structural work focused on the static IMiD– CRBN binary complex and neosubstrate–IMiD–CRBN ternary complex has revealed that IMiDs directly bind to the CRBN thalidomide-binding domain (TBD) and create a neo-interface to engage substrate degrons, such as the G-loop motif, to enable subsequent ubiquitylation and degradation of the target protein ^16-33^. Recent advances have further revealed dynamic allostery: IMiDs trigger CRBN conformational changes from an “open” to a “closed” state, facilitating the binding of Ikaros transcription factors ^24^. Although these studies have advanced our understanding of how IMiDs glue the neosubstrate and CRBN ^12-15^, the complete molecular picture of how IMiDs glue the neosubstrate to induce ubiquitylation within the intact, catalytically active CRL4^CRBN^ ligase complex remains unknown, which is of particular importance in the context that the optimization of IMiD pharmacology based on the ternary complex models does not fully correlate with therapeutic efficacy ^19,34^.

Here, we present cryo-electron microscopy (cryo-EM) structures of eight approved and clinical IKZF3-targeting IMiDs in complex with the neosubstrate IKZF3 and fully Nedd8-activated CRL4^CRBN^ (_N8_CRL4^CRBN^) ligase and provide the first comprehensive molecular portrait of how IMiDs engage the intact, catalytically active CRL4^CRBN^ ubiquitylation machinery to induce neosubstrate ubiquitylation. We unexpectedly observed that “next-generation” IMiDs, such as mezigdomide and golcadomide, can additionally bind to the non-degron zinc finger 3 (ZF3) of IKZF3 via an unprecedented cryptic gluing-driven interfacial (GDI) pocket, mediating additional drug–protein interactions and enhancing IKZF3 ubiquitylation and degradation efficiency. Thus, the full molecular portrait of IMiD-induced neosubstrate ubiquitylation offers more comprehensive and functionally relevant mechanistic insights into the efficacy of IMiDs than binary or ternary complexes do, informing a structure-guided design strategy targeting the GDI pocket to improve therapeutic efficacy.

### Approved and clinical IMiDs induce IKZF3 ubiquitylation and mediate IKZF3 degradation with varying efficiencies

Currently, eight approved or clinical IMiDs with known chemical structures have been reported to degrade IKZF3 (Fig. S1A): thalidomide ^7^, lenalidomide (CC-5013) ^7-9^, pomalidomide (CC-4047) ^7-9^, avadomide (CC-122) ^10,35^, iberdomide (CC-220) ^29,36^, mezigdomide (CC-92480) ^37^, golcadomide (CC-99282) ^38,39^, and cemsidomide (CFT7455) ^40^. Prior proteomic profiling of MM1S multiple myeloma cells identified lysine 166 (K166) as the top-ranking IMiD-regulated ubiquitylation site located within the zinc finger 2 (ZF2) degron motif of IKZF3 ^7^. In our experiments, we first verified the potency of these eight IMiDs to induce IKZF3 degradation in cells (Figs. S1B, S1C), providing a relative ranking of the IKZF3 degradation-inducing potency of these eight IMiDs: thalidomide and lenalidomide exhibited weaker potency; pomalidomide and avadomide showed enhanced potency; and iberdomide, mezigdomide, golcadomide, and cemsidomide demonstrated strong IKZF3 degradation-inducing potency (Figs. S1B, S1C). Next, we conducted mass spectrometry analysis of IKZF3 ubiquitylated *in vitro* in the presence of IMiDs (Figs. S1D, S1E). Three IKZF3 lysine residues (K158, K166, and K172) modified with the GG remnant were consistently identified in eight IMiD-induced ubiquitylation reactions (Fig. S1D). Among them, K166 had the highest number of peptide-spectrum matches (PSMs), confirming that it is the most extensively ubiquitylated lysine (Fig. S1D). Taken together, these data show that eight IMiDs can induce IKZF3 degradation with varying efficiencies and that the lysine residue K166 of IKZF3 is the preferential ubiquitylation site.

### Cryo-EM structures unveil a unified conformation of eight _N8_CRL4^CRBN^–IMiD–IKZF3 active ubiquitylation assemblies

Next, we aimed to determine the cryo-EM structures of a fully assembled active _N8_CRL4^CRBN^ ligase with the UbcH5a∼Ub thioester (‘∼’ represents an activated thioester bond between the donor ubiquitin and the E2 catalytic cysteine), positioned at the catalytic apex for ubiquitin transfer to IKZF3 K166 in the presence of eight discrete IMiDs. Since the IMiD-induced substrate ubiquitylation transition state is fleeting (Fig. S2A), an activity-based chemical trapping strategy ^41-43^ was employed to engineer covalent connectivity between the donor ubiquitin carboxyl terminus and the IKZF3 K166 site (Figs. S2B–S2D) and crosslink to the UbcH5a catalytic cysteine, facilitating the formation of a stable topological mimic of the native ubiquitylation transition state (Fig. S2A). Eight complexes were reconstituted in the presence of individual IMiD molecules and purified via size-exclusion chromatography (SEC, Fig. S2E and 2F). Single-particle cryo-EM analysis yielded high-quality maps of eight _N8_CRL4^CRBN^–IMiD–IKZF3 ubiquitylation assemblies, which were refined to an overall resolution of 3.4–3.8 Å (Figs. 1A and 1B, Fig S3, table S1 and S2, and data S1). The ubiquitylation assembly adopted a large, closed, carabiner-like conformation (180 Å by 120 Å, Fig. 1B), with all expected individual components clearly observed and unambiguously modeled (Figs. 1B and 1C, Fig. S4), including the IMiD molecules (Figs. 1B, 1C).

**Figure. 1.**
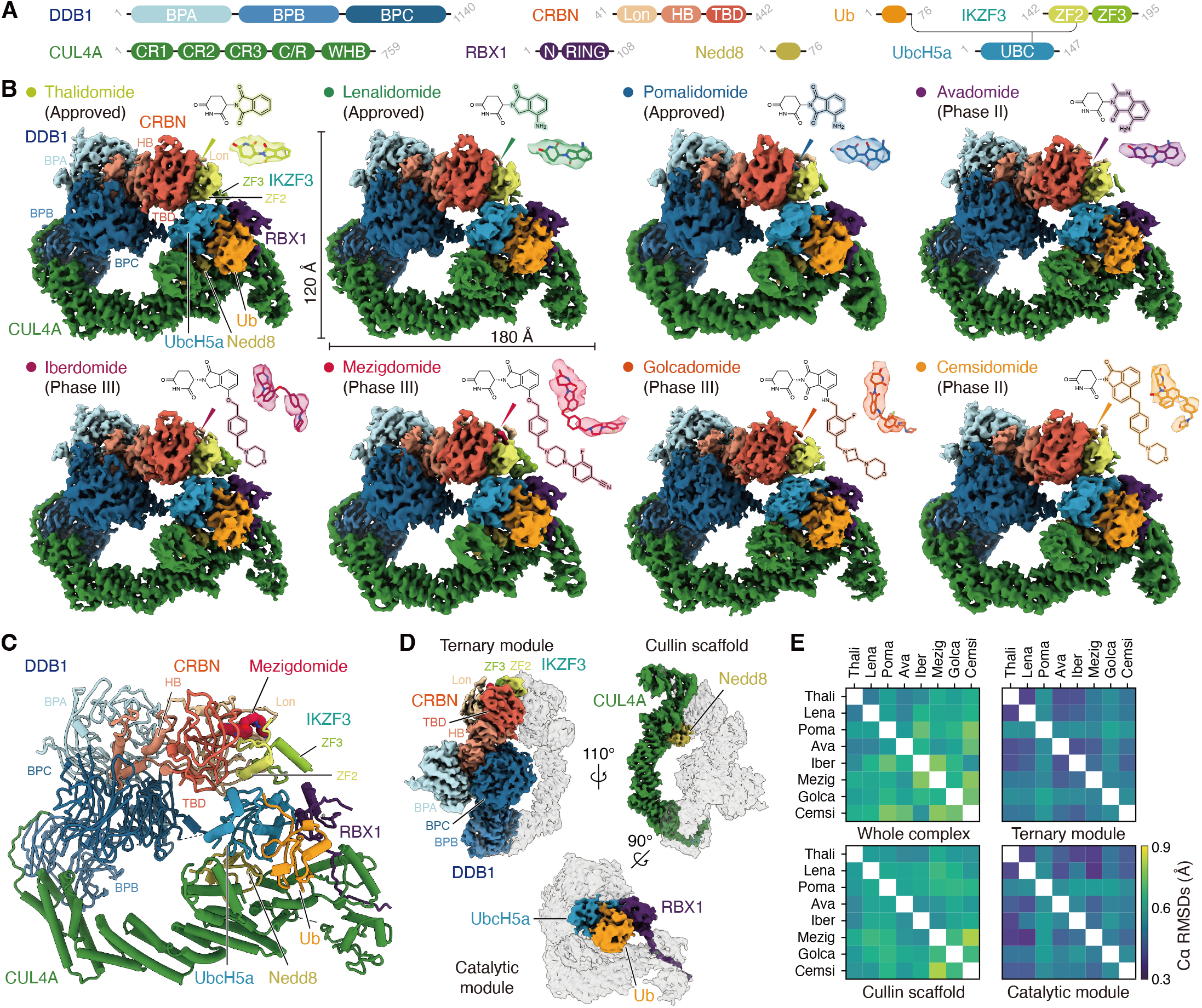
Structural visualization of _N8_CRL4^CRBN^ E3 ligase-mediated IKZF3 ubiquitylation induced by eight approved and clinical IMiDs. (**A**) Colored domain scheme of subunits with the catalytically active CRL4^CRBN^ ubiquitylation assembly. The chemical crosslink between Ub C-terminus, IKZF3 K166, and UbcH5a active site is indicated by lines. (**B**) Cryo-EM maps representing eight IKZF3-targeting IMiDs in complex with IKZF3 and E2∼Ub loaded _N8_CRL4^CRBN^ ligase with the active ubiquitylation assembly. Chemical structures, cartoon models, and cryo-EM densities of eight individual IMiDs are shown. (**C**) Cartoon model of the mezigdomide-organized ubiquitylation assembly with subcomponents indicated. (**D**) Architectures of the ternary module, the cullin scaffold, and the catalytic module in the mezigdomide-organized ubiquitylation assembly. (**E**) Heat map graphs showing Cα RMSDs of the whole complex, the ternary module, the cullin scaffold, and the catalytic module by pairwise alignments of eight structures.

The intact ubiquitylation assembly was structurally divided into three functional modules: the ternary module, the cullin scaffold, and the catalytic module (Fig. 1D). The ternary module consists of the substrate adaptor DDB1, the substrate receptor CRBN, the target protein IKZF3, and the IMiD molecule sandwiched by CRBN and IKZF3. The intermedial helical bundle (HB) of CRBN is embraced by two DDB1 β-propeller folds, BPA and BPC (Fig. 1D). The phthalimide/glutarimide cores of eight individual IMiDs are deeply buried in the CRBN thalidomide-binding domain (TBD) pocket, creating a neo-interface for engaging the G-loop of the IKZF3 ZF2 motif (Fig. S5A). In the cullin scaffold, CUL4A adopts an arc-shaped conformation, with its activator Nedd8 covalently modified on the WHB domain (Fig. 1D). The catalytic module contains a RING-type E3 ligase, RBX1, and UbcH5a∼Ub (Fig. 1D). RBX1 binds UbcH5a through a canonical RING domain–E2 interface, embracing and activating the ubiquitin in a “closed” conformation (Fig. 1D, Fig. S5B) ^41,42,44-46^. The ternary module, the cullin scaffold, and the catalytic module are interconnected through densely packed interactions (Fig. 1D, Fig S5B). For example, (1) the CUL4A N-terminal extension sequence and CR1 domain of CUL4A wrap around the second DDB1 β-propeller fold, BPB, via electrostatic and hydrophobic interactions ^47^, connecting the ternary module and the cullin scaffold (Fig. S5B); (2) the C-terminal C/R domain of CUL4A binds RBX1 and the Nedd8 modified on the WHB domain interacts with the UbcH5a backside ^44^, positioning the RBX1/UbcH5a∼Ub catalytic module to juxtapose with the substrate IKZF3 in the IMiD-organized ternary module (Fig. 1D, Fig S5B).

Pairwise comparisons of the whole complex of these eight ubiquitylation assemblies, each consisting of eight protein subunits and approximately 2,600 amino acid residues, revealed Cα root mean square deviation (RMSD) values within a remarkably narrow range, from 0.3 to 0.9 Å (Fig. 1E, Fig. S6A). Alignments of the ternary module, the cullin scaffold, or the catalytic module also yielded similarly low Cα RMSD values (Fig. 1E). These results indicate that the structures of the _N8_CRL4^CRBN^–IMiD–IKZF3 ubiquitylation assemblies exhibit a high degree of conformational conservation, despite the structural differences among the IMiD molecules.

### IMiDs-dependent ternary complex is tightly packed into active ubiquitination assemblies through conformational plasticity of the _N8_CRL4^CRBN^ ligase

In eight active ubiquitylation assemblies, we observed a consistently multi-layered, close packing in the IMiD binding region and the substrate ubiquitylation region (Fig. 2A). In the IMiD binding region, all eight IMiD molecules simultaneously bind to both the CRBN TBD domain and the IKZF3 ZF2 degron through the phthalimide/glutarimide cores (Figs. 2B, 2C), juxtaposing the K166 of IKZF3, the UbcH5a catalytic cysteine, and the ubiquitin C-terminus in the substrate ubiquitylation region (Fig. S6B). Notably, in all eight structures, the spatial distances between the Cα atoms of these residues (IKZF3 K166, UbcH5a C85, and Ub G75) meet the requirements for the chemical bonds and atomic arrangement in the native transition state intermediate of the ubiquitin transfer reaction (Figs. S6C, S6D).

**Figure. 2.**
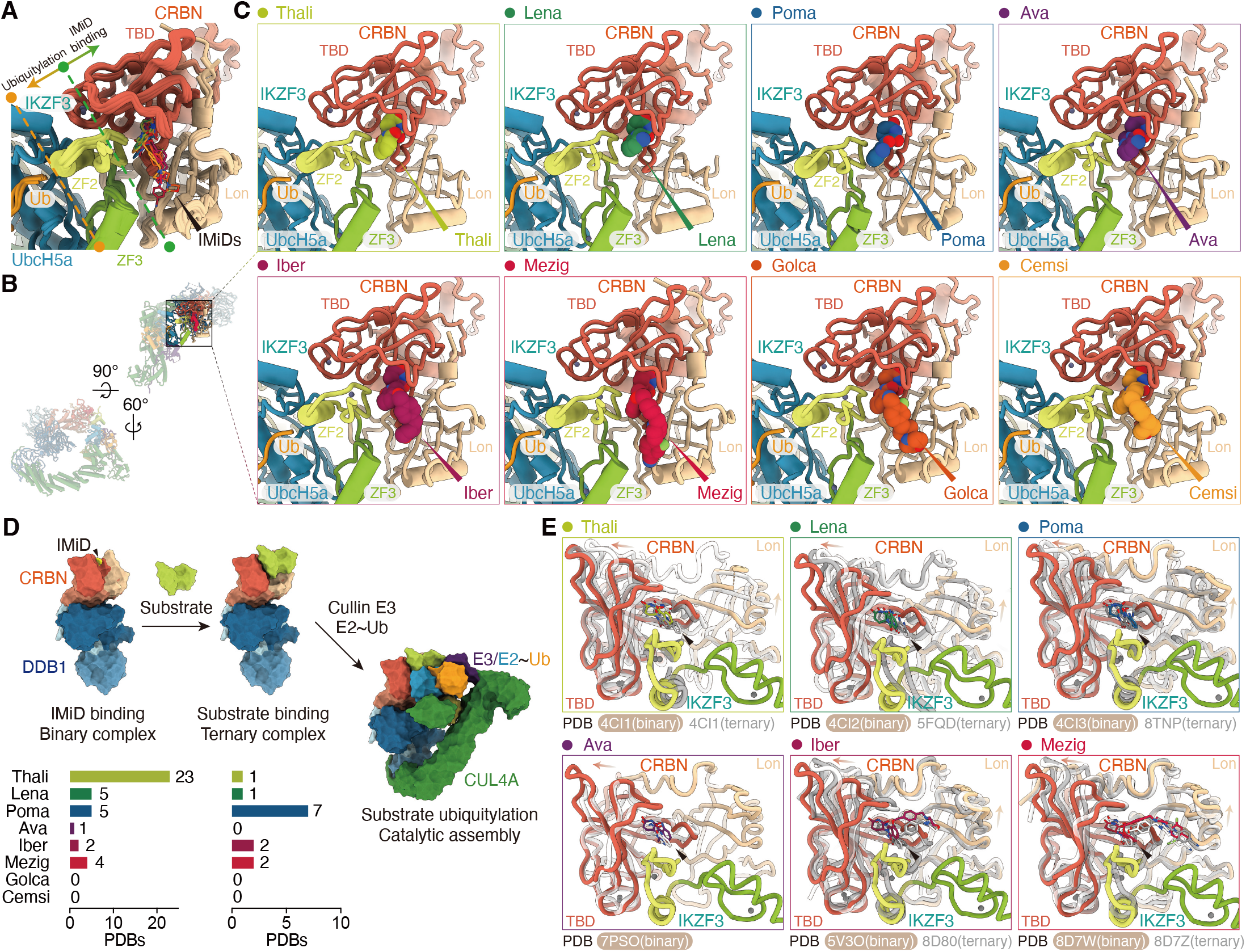
Conserved binding mode of eight IMiDs in inducing neosubstrate ubiquitylation. (**A**) A close-up view of the IMiD binding region and substrate ubiquitylation region in eight IMiD-organized ubiquitylation assemblies. (**B**) Guide to viewing the IMiD binding regions and substrate ubiquitylation regions shown in **C**. (**C**) Close-up views showing the binding mode of eight IMiDs. (**D**) Cartoon diagram illustrating the pathway for IMiD-organized complex assembly and summary of reported PDB structures of binary or ternary complexes containing eight IMiDs studied here. (**E**) Structural alignments of individual IMiD-organized binary complex, ternary complex, and ubiquitylation assembly, highlighting the conformational changes of CRBN TBD and Lon domains.

The CUL4A-interacting β-propeller BPB of DDB1, previously reported to adopt “hinged”, “linear”, or “twisted” conformations in apo CRBN/DDB1 complexes ^24^, consistently adopts the “hinged” conformation in all eight structures (Fig. S7A). A comparison of the CRBN conformations in our _N8_CRL4A^CRBN^–Mezigmezigdomide–IKZF3 complex (hereafter, the Mezig complex) and an IKZF1–mezigdomide–CRBN/DDB1 ternary complex (PDB 8D7Z) ^24^ revealed that while CRBN maintains its DDB1-buried HB domain almost unchanged, the IMiD-binding TBD and Lon domains undergo co-directional movements (Fig. S7B). Structural comparisons between previously reported binary/ternary complexes (Fig. 2D) and the ubiquitylation complexes of six IMiDs (excluding golcadomide and cemsidomide, for which no structural data of binary/ternary complexes are available) reveal that both the TBD and Lon domains of CRBN undergo stepwise displacements (Fig. 2E), reflecting their role in coordinating substrate positioning during ubiquitylation.

As a multi-subunit E3 ligase complex, CRL4A E3 ligase exhibits a modular architecture that allows it to adopt distinct states ^48^. For example, the CRL4A^DDB1/SV5V^ complex (PDB 2HYE) ^47^ represents an inactive state lacking Nedd8, E2∼Ub, and substrate; the CRL4A^DDB1/DDB2^–DNA complex (PDB 4A0K) ^49^ binds a substrate-related DNA-duplex; and the Nedd8-activated _N8_CRL4A^DDB1/CSA^–UVSSA–E2–Ub complex (PDB 8B3G) ^45^ is fully loaded with E2∼Ub and substrate (Fig. S7C). The alignment of these structures with our ubiquitylation catalytic intermediate-mimicking Mezig complex reveals substantial conformational flexibility of stepwise rotations or displacements of the DDB1 BPA domain (Fig. S7D, S7F), the RBX1/E2∼Ub catalytic module (Fig. S7E, S7G), the WHB domain and the C/R domain of CUL4A (Figs. S7H, 7I). These conformational changes match the steps of substrate binding (PDB 4A0K), Nedd8 activation/E2∼Ub loading (PDB 8B3G), and ubiquitin transfer (the Mezig structure), starting from the CRL4A^DDB1/SV5-V^ complex (PDB 2HYE, Figs. S7C–S7E).

### “Next-generation” IMiDs bind a cryptic gluing-driven interfacial (GDI) pocket formed by the ZF3 motif and the CRBN Lon domain

Structural analysis of the eight complexes unexpectedly revealed that four IMiDs, iberdomide, mezigdomide, golcadomide, and cemsidomide, which exhibit strong IKZF3 degradation potency, engage both the degron motif ZF2 and the non-degron motif ZF3 of IKZF3 (Fig. 3A, Fig. S8). The refined cryo-EM densities supported the atomic modeling of the IKZF3 non-degron motif ZF3 (Fig. 3B), which had not been observed in previous structures of the ternary complex containing mezigdomide or iberdomide (Fig. 3C) ^24^. The IKZF3 non-degron motif ZF3, which serves as a “proteinaceous glue”, fills the groove near the IMiD binding region and the substrate ubiquitylation region formed by the CRBN Lon domain and RBX1 (Fig. 3A, Fig. S8). The involvement of IKZF3 ZF3 in contact with the CRBN Lon domain echoes the previous observation that the deletion of ZF3 from the ZF2–ZF3 construct results in a tenfold decrease in binding affinity for the IMiD–CRBN complex ^28^. Interestingly, a previously reported ternary complex structure (PDB 8DEY) ^23^, which contains another Ikaros family transcription factor IKZF2, whose tandem ZF2–ZF3 domains share 91.2% sequence identity with those of IKZF3 (Fig. S1F), has revealed an alternative conformation of ZF3 (Fig. S8B). Superposition of the IKZF2 ternary complex with our Mezig structure revealed an ∼100°angular difference between the ZF3 motifs of IKZF2 and IKZF3. When modeled into the mezigdomide-organized ubiquitylation assembly, the IKZF2 ZF3 would clash with UbcH5a (Fig. S8B). These findings demonstrate that resolving the full CRL4^CRBN^ ubiquitylation complex is essential for accurately capturing critical neosubstrate features, such as the ZF3 motif, which could otherwise be obscured or misrepresented in ternary complexes.

**Figure. 3.**
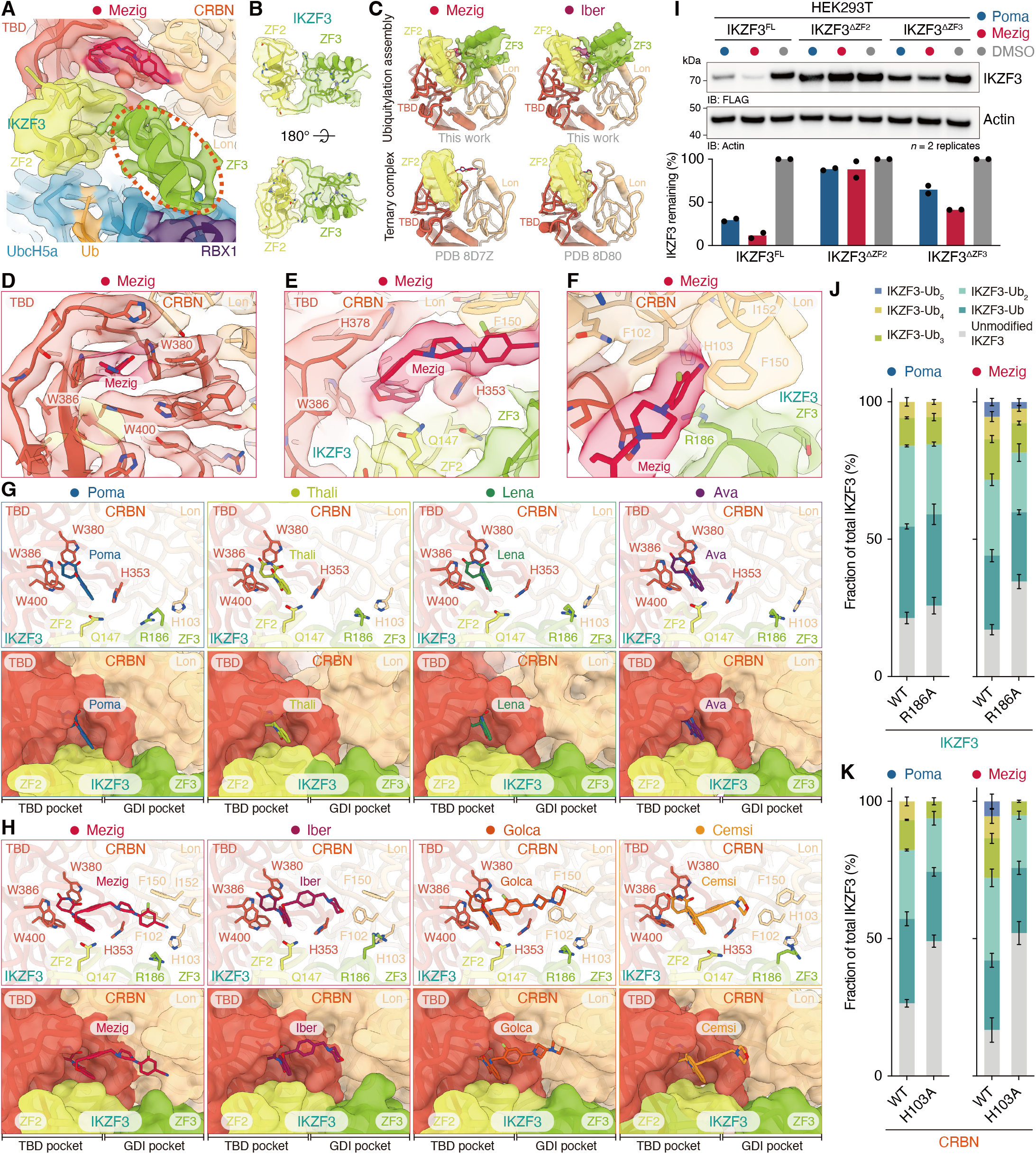
A cryptic GDI pocket formed by the IKZF3 non-degron motif ZF3 and CRBN Lon domain contributes to IMiD-induced IKZF3 ubiquitylation and degradation. (**A**) A close-up view of the Mezig structure with a cartoon model fit into the cryo-EM map. The red dashed circle indicates the unexpected IKZF3 ZF3 density. (**B**) Two orientations of the IKZF3 cryo-EM density in the Mezig structure with cartoon model fit. (**C**) Comparison of tandem ZF2–ZF3 motifs observed in mezigdomide- or iberdomide-organized ubiquitylation assemblies (top, this work) and ternary complexes (PDB 8D7Z, 8D80), revealing the absence of ZF3 density in ternary complexes. The ZF2–ZF3 motifs in ubiquitylation assemblies are shown as semi-transparent cryo-EM densities with cartoon model fit. The ZF2 motifs in ternary complexes (PDB 8D7Z, 8D80) are shown as semi-transparent cartoon surfaces and models. (**D**) Detailed view of mezigdomide binding in the CRBN TBD pocket. Key TBD pocket residues, CRBN W380, W386, and W400, are indicated. (**E**) Detailed view of mezigdomide interacting with CRBN and IKZF3. Key interacting residues, CRBN F150, H353, H378, W386, and IKZF3 Q147 are indicated. (**F**) Detailed view of mezigdomide interacting with the IKZF3–CRBN GDI pocket. Key interacting residues, CRBN F102, H103, F150, I152, and IKZF3 R186, are indicated. Cryo-EM density map and atomic model are shown. (**G, H**) Zoomed-in views of each IMiD molecule binding in the CRBN TBD pocket or GDI pocket. Side chains of key interacting amino acid residues are shown as stick models (top). Surface representations of CRBN and IKZF3 are used to highlight the structural features of the binding pockets. (**I**) A representative western blot image (top) of *in vivo* degradation assay using IKZF3^FL^, IKZF3^ΔZF2^, or IKZF3^ΔZF3^ construct and bar graph showing quantified IKZF3 degradation data from *n* = 2 technical replicates (bottom). (**J**) *In vitro* ubiquitylation assay using WT IKZF3 ZF2–ZF3 or its R186A mutant in the presence of pomalidomide or mezigdomide. (**K**) *In vitro* ubiquitylation assay using WT CRBN or its H103 mutant with the fully-assembled _N8_CRL4^CRBN^ E3 ligase complex. Bar graph comparing the fraction of unmodified and ubiquitylated IKZF3 species in the presence of eight IMiDs. Data are presented as mean ± SD from *n* = 3 technical replicates.

In the ubiquitylation assembly, the ZF3 motif and the CRBN Lon domain enclose a pocket at the protein–protein interface (Fig. 3A). In contrast to the canonical TBD binding pocket, this distinct interfacial pocket only becomes apparent only upon the IMiD-induced binding of IKZF3 to CRBN during the substrate ubiquitylation process. We thus defined this cryptic pocket as a gluing-driven interfacial (GDI) pocket. In all eight IMiD-organized ubiquitylation assemblies, the phthalimide/glutarimide cores of IMiDs conservatively bind to the canonical CRBN TBD pocket, making contacts with the CRBN aromatic residues H353, W380, W386, and W400 (Figs. 3D, 3G, 3H). The only side-chain interaction between IKZF3 ZF2 and IMiDs is formed by IKZF3 Q147 (Figs. 3E, 3G, 3H), whose substitution with histidine (Q147H) results in IMiD resistance ^7^. Intriguingly, the additional groups of the “next-generation” IMiDs (mezigdomide, iberdomide, golcadomide, and cemsidomide) reach the GDI pocket juxtaposed with the TBD pocket, making contacts with IKZF3 R186 and hydrophobic amino acid residues (e.g., H103, F102, and F150) on the CRBN Lon domain (Figs. 3F, 3G, 3H). Based on their binding modes with the TBD or GDI pockets, the eight IKZF3-targeting IMiDs can be classified into two categories: monovalent IMiDs (MvIMiDs, Fig. S9A), which bind exclusively to the TBD pocket (e.g., pomalidomide, thalidomide, lenalidomide, and avadomide), and bridging IMiDs (BdIMiDs, Fig. S9B), which simultaneously engage both the TBD and GDI pockets to introduce additional drug–protein interactions (e.g., mezigdomide, iberdomide, golcadomide, and cemsidomide).

We performed IKZF3 degradation assays in cells and mutagenesis-based *in vitro* ubiquitylation assays to validate our structural observation that the IKZF3 ZF3 motif contributes to GDI pocket formation, and to investigate how the IMiD–GDI pocket interaction influences IKZF3 ubiquitylation and degradation. HEK293T cells transfected with IKZF3 (IKZF3^FL^ construct) or IKZF3 deletion constructs were treated with 1 µM of either pomalidomide or mezigdomide for 8 hours, and the intracellular abundance of IKZF3 was assessed by western blot (WB, Fig. 3I). The deletion of the IKZF3 degron motif ZF2 (amino acid residues 142–169) (IKZF3^ΔZF2^ construct) rendered IKZF3 resistant to IMiD treatment, effectively serving as a reliable negative control (Fig. 3I). Notably, the deletion of the IKZF3 non-degron motif ZF3 (amino acid residues 171–198) (IKZF3^ΔZF3^ construct) also resulted in IMiD resistance, although to a lesser extent than the IKZF3^ΔZF2^ construct did (Fig. 3I). These *in vivo* results agree with the findings from previous cellular IKZF1/3 zinc finger deletion degradation assays that deletion of ZF3 impaired IMiD-induced IKZF3 degradation using fluorescent reporter vectors ^28^. The substitution of the IKZF3 residue R186 with alanine (R186A) impaired IKZF3 ubiquitylation induced by mezigdomide *in vitro* (Fig. 3J), suggesting that the R186A mutation disrupts additional drug–protein interactions within the GDI pocket. Our *in vitro* data were further substantiated by previous results that the mutation of this consensus arginine residue in Ikaros family transcription factors (IKZF3 R186 and IKZF1 R185) impairs the binding affinity of ZF2–ZF3 construct to the IMiD–CRBN complex ^28^. The Lon domain side of the GDI pocket centers on residue H103 (Fig. S10A) and includes a hydrophobic surface (Figs. S10B, S10C) that synergistically recruits MGDs and zinc finger proteins (Fig. S10D) ^18,22^. The incorporation of the CRBN H103A mutant into the _N8_CRL4^CRBN^ ligase complex impaired IMiD-induced IKZF3 ubiquitylation (Fig. 3K).

Together, our structural and biochemical findings reveal that the IKZF3 non-degron motif ZF3 glues the CRBN Lon domain and RBX1/UbcH5a catalytic module to form a cryptic GDI pocket and that the IMiD–GDI pocket interactions provide an unexpected mechanistic rationale for the enhanced efficacy of BdIMiDs compared to that of MvIMiDs.

### Understanding IMiD efficacy through IMiD–GDI pocket interactions in the full CRL4^CRBN^ ubiquitylation complex

Although IMiDs all share phthalimide/glutarimide core structures, different IMiDs have been shown to reshape the CRBN to selectively recognize distinct proteins, such as IKZF3, GSPT1, and CK1α ^12^. We conducted *in silico* docking of a panel of previously reported IMiDs with distinct substrate selectivities ^19,27,30,31,50-55^ into the fully assembled CRL4^CRBN^ ubiquitylation complex using the Schrödinger Maestro software (Fig. 4A, Fig. S11). The analysis revealed that: (1) all eight IKZF3-targeting IMiDs characterized in this study exhibited relatively low (and thus more favorable) docking scores (Fig. 4A); (2) the four BdIMiDs with enhanced IKZF3 selectivity and degradation potency showed significantly lower scores than the MvMiDs, with mezigdomide achieving the most favorable score (Fig. 4A); and (3) IMiDs selective for other neosubstrates such as GSPT1 or CK1α, as well as promiscuous degraders, generally yielded relatively unfavorable docking scores (Fig. 4A). However, performing the same *in silico* analysis using the previously reported ternary complex (PDB 8D7Z) failed to recapitulate these results (Fig. 4A).

**Figure. 4.**
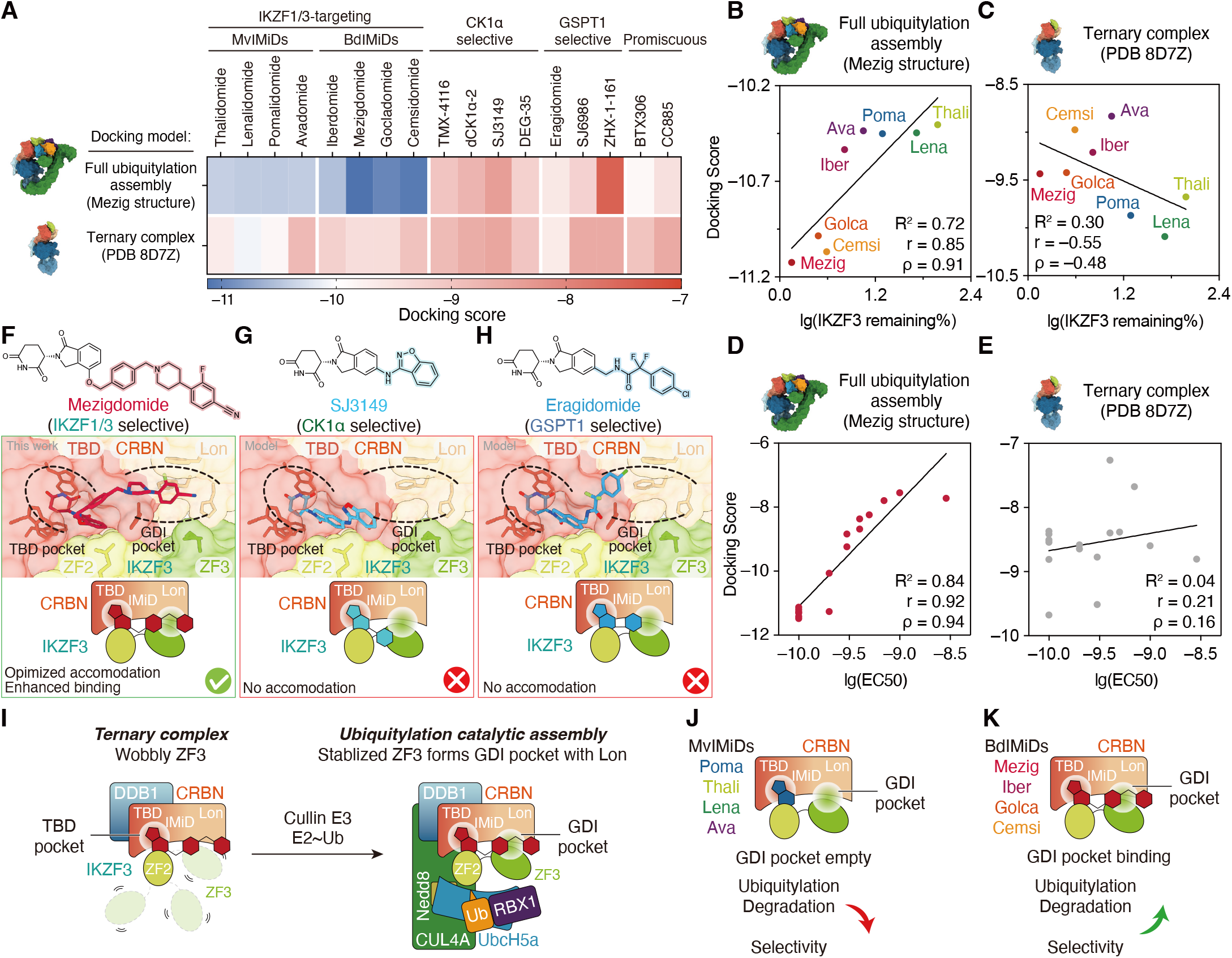
The IMiD–GDI pocket interplay in the substrate ubiquitylation process provides a rational understanding of IMiDs’ efficacy. (**A**) Heat map graphs showing docking scores of IMiDs with distinct target selectivities. mezigdomide-organized full ubiquitylation assembly (Mezig complex) or ternary complex (PDB 8D7Z) was used as the docking model. Low (favorable) and high (unfavorable) docking scores are indicated by blue and red colors. (**B, C**) Dot plots showing the correlation analysis of the logarithm of IKZF3 degradation potency [lg(IKZF3 remaining%)] for the eight IKZF3-targeting IMiDs against their docking scores within the full ubiquitylation complex (**B**) or ternary complex (**C**). IKZF3 degradation potency data are from **Fig. S1B**. (**D, E**) Dot plots showing the correlation analysis of the logarithm of previously reported IKZF3 EC50 [lg(EC50)] for mezigdomide analogs against their docking scores within the full ubiquitylation complex (**D**) or ternary complex (**E**). R^2^ of the simple linear regression model, Pearson’s correlation coefficient (r), and Spearman’s correlation coefficient (ρ) are indicated. (**F**) Structural analysis and schematic illustration of mezigdomide engaging both the CRBN TBD and GDI pockets in our cryo-EM structure of full ubiquitylation assembly. (**G, H**) Structural analysis and schematic illustration of GSPT1-selective eragidomide (**G**) or CK1α-selective SJ3149 (**H**) docked into our Meizg structure. (**I**) Cartoon diagram illustrating the pathway for the IMiD-organized ternary complex to catalytically active ubiquitylation assembly. Engagement of the fully-assembled _N8_CRL4^CRBN^ E3 ligase and E2∼Ub enables the wobbly IKZF3 ZF3 motif to form the GDI pocket. (**J, K**) Cartoon summary illustrating the rationale for understanding the molecular origin of the efficacy of MvIMiDs (**J**) or BdIMiDs (**K**) through IMiD–GDI pocket interplay.

The therapeutic efficacy of IMiDs encompasses both substrate selectivity and degradation potency. We performed a correlation analysis by plotting the logarithm of IKZF3 degradation potency for the eight IKZF3-targeting IMiDs against their docking scores. Using the CRL4^CRBN^ full ubiquitylation complex, linear regression yielded an R^2^ of 0.74, with a Pearson’s correlation coefficient (r) of 0.85 and a Spearman’s correlation coefficient (ρ) of 0.91 (Fig. 4B). By contrast, the ternary complex model (PDB 8D7Z) resulted a markedly lower R^2^ of 0.30, with negative r = – 0.55 and ρ = –0.48 (Fig. 4C). We further strengthened the application of our discovery of the IMiD–GDI pocket interaction in the full ubiquitylation complex for understanding the structure-activity relationship (SAR) of mezigdomide analogs. Based on SAR data from the mezigdomide development study ^37^, docking scores from the full ubiquitylation complex showed a strong positive correlation with IKZF3 degradation potency (Fig. 4D), while in contrast, docking scores from the ternary complex model (PDB 8D7Z) showed a poor correlation (Fig. 4E). These molecular docking analyses revealed that enhanced drug–GDI pocket compatibility correlates with improved efficacy (Fig. S12), suggesting a previously unrecognized strategy for the SAR optimization of IMiDs.

We further selected four representative IMiDs with distinct substrate selectivities for structural analysis. In our resolved Mezig structure, the GDI pocket formed by the IKZF3 ZF3 motif and CRBN Lon domain is optimally accommodated by the terminal 2-fluoro-4-cyanophenyl group of mezigdomide, establishing extra drug–protein interactions (Fig. 4F). However, in the docking models, the GSPT1-selective degrader eragidomide ^27^ (Fig. 4G) or the CK1α-selective degrader SJ3149 ^19^ (Fig. 4H) could not be accommodated within the GDI pocket. These analyses indicate that steric or chemical incompatibility with the GDI pocket can hinder effective IMiD engagement in the full ubiquitylation machinery, thereby influencing substrate selectivity.

Collectively, our results show that *in silico* docking with the full CRL4^CRBN^ ubiquitylation complex enables clear discrimination of IKZF3-targeting IMiDs and elucidation of SAR via the IMiD–GDI pocket interplay, insights unattainable from the ternary complexes.

## Discussion

Thalidomide, a historic milestone in medical science, was initially notorious for its severe teratogenic effects ^1^ but was subsequently repurposed as an effective therapy for multiple myeloma ^56^. Thalidomide-derived IMiDs are now considered the most successful targeted protein degradation drugs both for clinical treatment and market application ^6^, exerting therapeutic effects by reprogramming the substrate selectivity of the CRL4^CRBN^ E3 ubiquitin ligase ^3-5^ and degrading disease-associated proteins, such as IKZF1 and IKZF3 ^7-9^. In the present study, leveraging activity-based trapping chemistry ^41-43^, we provide the first comprehensive molecular portrait of how IMiDs engage the intact, catalytically active CRL4^CRBN^ ubiquitylation machinery to induce neosubstrate ubiquitylation, thereby advancing beyond previous mechanistic insights gained from partial ternary complexes.

Our cryo-EM structures of eight catalytically active _N8_CRL4A^CRBN^–IKZF3–E2–Ub complexes induced by distinct IMiDs reveal a conserved architecture of the ubiquitylation machinery driving neosubstrate modification. Structural and functional analyses revealed that the non-degron ZF3 motif of IKZF3 plays an unexpected regulatory role, acting as a “proteinaceous glue” to reinforce the compact architecture of the ubiquitylation region and enhance IMiD-induced ubiquitylation efficiency. Compared to ternary complexes lacking the activated _N8_CRL4^CRBN^ E3 ligase or E2∼Ub conjugates, where ZF3 is either unresolved ^24^ or differently positioned ^23^, our fully assembled ubiquitylation complexes reveal that ZF3 adopts a distinct conformation, positioned between the CRBN Lon domain and the RBX1/E2∼Ub catalytic module (Fig. 4I). This arrangement enables the formation of a previously unrecognized GDI pocket that enhances IMiD binding (Fig. 4I). Relative to MvIMiDs (e.g., pomalidomide, thalidomide, lenalidomide, and avadomide), which bind only the CRBN TBD pocket (Fig. 4J), BdIMiDs (e.g., mezigdomide, iberdomide, golcadomide, and cemsidomide) engage both TBD and GDI pockets (Fig. 4K), forming additional interactions that enhance degradation potency and target selectivity. Furthermore, *in silico* docking using the fully assembled CRL4^CRBN^ ubiquitylation complex enables precise differentiation of IKZF3-targeting IMiDs and facilitates SAR interpretation through the newly uncovered IMiD– GDI pocket interactions, which are not achievable with previous ternary complex models. Together, these findings highlight the importance of resolving the full ubiquitylation machinery structure to provide a more comprehensive mechanistic basis for understanding IMiDs’ efficacy and selectivity.

To date, the vast majority of CRBN-based molecular glue degraders (MGDs) exploit the canonical and conserved phthalimide/glutarimide–TBD pocket interaction ^16-33^, restricting the chemical space available for modulating substrate selectivity or minimizing off-target effects through structure-based molecular design. Our findings reveal a comprehensive structural view of the IMiD-organized, fully assembled ubiquitylation machinery and open the possibility of designing improved MGDs that are tailored to engage the GDI pocket, thereby enabling more efficient and selective targeted substrate ubiquitylation through more tightly organized and catalytically competent ubiquitylation machinery. We anticipate that the newly uncovered interaction between BdIMiDs and the GDI pocket may pave the way for a new paradigm in the structure-guided rational design of improved MGDs.

## Acknowledgments

We acknowledge the Tsinghua University Branch of China National Center for Protein Sciences (Beijing) and Shuimu BioSciences (Hangzhou) for cryo-EM data screening and collection. We thank the Center of Protein Analysis Technology, Tsinghua University, for MS analysis. We thank the National Natural Science Foundation of China (92253302, T2488301, 22137005, and 22227810 to L.L.; 22277073 and 92253302 to M.P.), National Key R&D Program of China (2022YFC3401500 to L.L.; 2023YFA0915300 to M.P.), and New Cornerstone Science Foundation (to L.L.). Shanghai Rising-Star Program (22QA1404900 to M.P.), Shanghai Pilot Program for Basic Research Shanghai Jiao Tong University (21TQ1400224 to M.P.), Foundation of Muyuan Laboratory (118602240 to M.P.). Fundamental Research Funds for the Central University (to M.P.). Shanghai Jiao Tong University 2030 Initiative (WH510363003/003 to M.P.), Shanghai Frontiers Science Center of Drug Target Identification and Delivery (ZXWH2170101 to H.A.). Z. D. thanks the support from Shuimu Tsinghua Scholars Program.

## Author contributions

Conceptualization: L.L., M.P., H.A., and Z.D. Chemical trapping methodology: Z.D. and Q.S. Cloning and protein purification: Z.D., Q.S., J.L., Z.H., L.Z., S.T., W.H., and Q.Z. Cryo-EM sample preparation: Z.D. Cryo-EM data processing: H.A. Model building: Z.D. and H.A. Ubiquitylation and degradation assays: Z.D., Q.S., and J.L. Molecular docking: Z.D., J.Z., and H.L. Data interpretation: Z.D. and H.A. Visualization: Z.D. Funding acquisition: L.L., M.P., and H.A. Supervision: L.L. Writing – original draft: Z.D., M.P., and H.A. Writing – review & editing: L.L., M.P., H.A., and Z.D.

## Competing interests

Authors declare that they have no competing interests.

## Data and materials availability

Coordinates and cryo-EM maps of IMiD-organized complexes have been deposited in the Protein Data Bank (PDB) and Electron Microscopy Data Bank (EMDB), including the thalidomide complex (PDB 9UUQ, EMD-64515), the lenalidomide complex (9V09, EMD-64658), the pomalidomide complex (9V0A, EMD-64659), the avadomide complex (9V0B, EMD-64660), the iberdomide complex (9V0C, EMD-64661), the mezigdomide complex (9UUM, EMD-64512), the golcadomide complex (9V0E, EMD-64662), and the cemsidomide complex (9V0F, EMD-64663). Newly created materials from this study may be requested from the corresponding authors.

## Notes

### Competing Interest Statement

The authors have declared no competing interest.

### Summary of Updates

We uploaded a new version and revised some typos

## References and Notes

1 McBride, W. G. THALIDOMIDE AND CONGENITAL ABNORMALITIES. The Lancet 278, 1358 (1961). 10.1016/S0140-6736(61)90927-8

2 Bartlett, J. B., Dredge, K. & Dalgleish, A. G. Timeline - The evolution of thalidomide and its IMiD derivatives as anticancer agents. Nat. Rev. Cancer 4, 314–322 (2004). 10.1038/nrc1323

3 Ito, T. et al./person-group>. Identification of a Primary Target of Thalidomide Teratogenicity. Science 327, 1345–1350 (2010). 10.1126/science.1177319

4 Lopez-Girona, A. et al./person-group>. Cereblon is a direct protein target for immunomodulatory and antiproliferative activities of lenalidomide and pomalidomide. Leukemia 26, 2326–2335 (2012). 10.1038/leu.2012.119

5 Jan, M., Sperling, A. S. & Ebert, B. L. Cancer therapies based on targeted protein degradation — lessons learned with lenalidomide. Nat. Rev. Clin. Oncol. 18, 401–417 (2021). 10.1038/s41571-021-00479-z

6 Miguel, J. S. et al./person-group>. Pomalidomide plus low-dose dexamethasone versus high-dose dexamethasone alone for patients with relapsed and refractory multiple myeloma (MM-003): a randomised, open-label, phase 3 trial. Lancet Oncol. 14, 1055–1066 (2013). 10.1016/S1470-2045(13)70380-2

7 Krönke, J. et al./person-group>. Lenalidomide Causes Selective Degradation of IKZF1 and IKZF3 in Multiple Myeloma Cells. Science 343, 301–305 (2014). 10.1126/science.1244851

8 Lu, G. et al./person-group>. The Myeloma Drug Lenalidomide Promotes the Cereblon-Dependent Destruction of Ikaros Proteins. Science 343, 305–309 (2014). 10.1126/science.1244917

9 Gandhi, A. K. et al./person-group>. Immunomodulatory agents lenalidomide and pomalidomide co-stimulate T cells by inducing degradation of T cell repressors Ikaros and Aiolos via modulation of the E3 ubiquitin ligase complex CRL4. Br. J. Haematol. 164, 811–821 (2014). 10.1111/bjh.12708

10 Krönke, J. et al./person-group>. Lenalidomide induces ubiquitination and degradation of CK1a in del(5q) MDS. Nature 523, 183–188 (2015). 10.1038/nature14610

11 List, A. et al./person-group>. Efficacy of lenalidomide in myelodysplastic syndromes. N. Engl. J. Med. 352, 549–557 (2005). DOI 10.1056/NEJMoa041668

12 Oleinikovas, V., Gainza, P., Ryckmans, T., Fasching, B. & Thomä, N. H. From Thalidomide to Rational Molecular Glue Design for Targeted Protein Degradation. Annu. Rev. Pharmacol. Toxicol. 64, 291–312 (2024). 10.1146/annurev-pharmtox-022123-104147

13 Konstantinidou, M. & Arkin, M. R. Molecular glues for protein-protein interactions Progressing toward a new dream. Cell Chem. Biol. 31, 1064–1088 (2024). 10.1016/j.chembiol.2024.04.002

14 Tsai, J. M., Nowak, R. P., Ebert, B. L. & Fischer, E. S. Targeted protein degradation: from mechanisms to clinic. Nat. Rev. Mol. Cell Biol. 25, 740–757 (2024). 10.1038/s41580-024-00729-9

15 Hughes, S. J. & Ciulli, A. Molecular recognition of ternary complexes: a new dimension in the structure-guided design of chemical degraders. Essays Biochem. 61, 505–516 (2017). 10.1042/Ebc20170041

16 Chrisochoidou, Y. et al./person-group>. Evaluating the impact of CRBN mutations on response to immunomodulatory drugs and novel CRBN-binding agents in myeloma. Blood (2025). 10.1182/blood.2024025861

17 Ting, P. Y. et al./person-group>. A molecular glue degrader of the WIZ transcription factor for fetal hemoglobin induction. Science 385, 91–99 (2024). 10.1126/science.adk6129

18 Mercer, J. A. M. et al./person-group>. Continuous evolution of compact protein degradation tags regulated by selective molecular glues. Science 383, eadk4422 (2024). doi:10.1126/science.adk4422

19 Nishiguchi, G. et al./person-group>. Selective CK1α degraders exert antiproliferative activity against a broad range of human cancer cell lines. Nat. Commun. 15, 482 (2024). 10.1038/s41467-024-44698-1

20 Kroupova, A. et al./person-group>. Design of a Cereblon construct for crystallographic and biophysical studies of protein degraders. Nat. Commun. 15, 8885 (2024). 10.1038/s41467-024-52871-9

21 Kerrigan, J. R. et al./person-group>. Discovery and Optimization of First-in-Class Molecular Glue Degraders of the WIZ Transcription Factor for Fetal Hemoglobin Induction to Treat Sickle Cell Disease. J. Med. Chem. 67, 20682–20694 (2024). 10.1021/acs.jmedchem.4c02251

22 Ma, X. et al./person-group>. Structural and biophysical comparisons of the pomalidomide- and CC-220-induced interactions of SALL4 with cereblon. Sci. Rep. 13, 22088 (2023). 10.1038/s41598-023-48606-3

23 Bonazzi, S. et al./person-group>. Discovery and characterization of a selective IKZF2 glue degrader for cancer immunotherapy. Cell Chem. Biol. 30, 235–247.e212 (2023). 10.1016/j.chembiol.2023.02.005

24 Watson, E. R. et al./person-group>. Molecular glue CELMoD compounds are regulators of cereblon conformation. Science 378, 549–553 (2022). doi:10.1126/science.add7574

25 Furihata, H. et al./person-group>. Structural bases of IMiD selectivity that emerges by 5-hydroxythalidomide. Nat. Commun. 11, 4578 (2020). 10.1038/s41467-020-18488-4

26 Matyskiela, M. E. et al./person-group>. Crystal structure of the SALL4-pomalidomide-cereblon-DDB1 complex. Nat. Struct. Mol. Biol. 27, 319–322 (2020). 10.1038/s41594-020-0405-9

27 Surka, C. et al./person-group>. CC-90009, a novel cereblon E3 ligase modulator, targets acute myeloid leukemia blasts and leukemia stem cells. Blood 137, 661–677 (2021). 10.1182/blood.2020008676

28 Sievers, Q. L. et al./person-group>. Defining the human C2H2 zinc finger degrome targeted by thalidomide analogs through CRBN. Science 362, eaat0572 (2018). doi:10.1126/science.aat0572

29 Matyskiela, M. E. et al./person-group>. A Cereblon Modulator (CC-220) with Improved Degradation of Ikaros and Aiolos. J. Med. Chem. 61, 535–542 (2018). 10.1021/acs.jmedchem.6b01921

30 Matyskiela, M. E. et al./person-group>. A novel cereblon modulator recruits GSPT1 to the CRL4(CRBN) ubiquitin ligase. Nature 535, 252–257 (2016). 10.1038/nature18611

31 Petzold, G., Fischer, E. S. & Thomä, N. H. Structural basis of lenalidomide-induced CK1α degradation by the CRL4CRBN ubiquitin ligase. Nature 532, 127–130 (2016). 10.1038/nature16979

32 Chamberlain, P. P. et al./person-group>. Structure of the human Cereblon-DDB1-lenalidomide complex reveals basis for responsiveness to thalidomide analogs. Nat. Struct. Mol. Biol. 21, 803–809 (2014). 10.1038/nsmb.2874

33 Fischer, E. S. et al./person-group>. Structure of the DDB1-CRBN E3 ubiquitin ligase in complex with thalidomide. Nature 512, 49–53 (2014). 10.1038/nature13527

34 Hassan, M. M. et al./person-group>. Exploration of the tunability of BRD4 degradation by DCAF16 trans-labelling covalent glues. Eur. J. Med. Chem. 279, 116904 (2024). 10.1016/j.ejmech.2024.116904

35 Hagner, P. R. et al./person-group>. CC-122, a pleiotropic pathway modifier, mimics an interferon response and has antitumor activity in DLBCL. Blood 126, 779–789 (2015). 10.1182/blood-2015-02-628669

36 Bjorklund, C. C. et al./person-group>. Iberdomide (CC-220) is a potent cereblon E3 ligase modulator with antitumor and immunostimulatory activities in lenalidomide- and pomalidomide-resistant multiple myeloma cells with dysregulated CRBN. Leukemia 34, 1197–1201 (2020). 10.1038/s41375-019-0620-8

37 Hansen, J. D. et al./person-group>. Discovery of CRBN E3 Ligase Modulator CC-92480 for the Treatment of Relapsed and Refractory Multiple Myeloma. J. Med. Chem. 63, 6648–6676 (2020). 10.1021/acs.jmedchem.9b01928

38 Carrancio, S. et al./person-group>. CC-99282 is a Novel Cereblon (CRBN) E3 Ligase Modulator (CELMoD) Agent with Enhanced Tumoricidal Activity in Preclinical Models of Lymphoma. Blood 138, 1200–1200 (2021). 10.1182/blood-2021-148068

39 Michot, J.-M. et al./person-group>. Clinical Activity of CC-99282, a Novel, Oral Small Molecule Cereblon E3 Ligase Modulator (CELMoD) Agent, in Patients (Pts) with Relapsed or Refractory Non-Hodgkin Lymphoma (R/R NHL) - First Results from a Phase 1, Open-Label Study. Blood 138, 3574–3574 (2021). 10.1182/blood-2021-147333

40 Perino, S. et al./person-group>. CFT7455, a Novel IKZF1/3 Degrader, Demonstrates Potent Anti-Tumor Activity in Models of Non-Hodgkin’s Lymphoma As a Single Agent or in Combination with Clinically Approved Agents. Blood 140, 11575–11575 (2022). 10.1182/blood-2022-166541

41 Li, J. et al./person-group>. Cullin-RING ligases employ geometrically optimized catalytic partners for substrate targeting. Mol. Cell 84, 1304–1320 e1316 (2024). 10.1016/j.molcel.2024.01.022

42 Liwocha, J. et al./person-group>. Mechanism of millisecond Lys48-linked poly-ubiquitin chain formation by cullin-RING ligases. Nat. Struct. Mol. Biol. 31, 378–389 (2024). 10.1038/s41594-023-01206-1

43 Davidson, G. A., Moafian, Z., Sensi, A. R. & Zhuang, Z. Thioether-mediated protein ubiquitination in constructing affinity- and activity-based ubiquitinated protein probes. Nat. Protoc. (2025). 10.1038/s41596-025-01162-8

44 Baek, K. et al./person-group>. NEDD8 nucleates a multivalent cullin-RING-UBE2D ubiquitin ligation assembly. Nature 578, 461–466 (2020). 10.1038/s41586-020-2000-y

45 Kokic, G. et al./person-group>. Structural basis for RNA polymerase II ubiquitylation and inactivation in transcription-coupled repair. Nat. Struct. Mol. Biol. 31, 536–547 (2024). 10.1038/s41594-023-01207-0

46 Cappadocia, L. & Lima, C. D. Ubiquitin-like Protein Conjugation: Structures, Chemistry, and Mechanism. Chem. Rev. 118, 889–918 (2018). 10.1021/acs.chemrev.6b00737

47 Angers, S. et al./person-group>. Molecular architecture and assembly of the DDB1-CUL4A ubiquitin ligase machinery. Nature 443, 590–593 (2006). 10.1038/nature05175

48 Petroski, M. D. & Deshaies, R. J. Function and regulation of cullin–RING ubiquitin ligases. Nat. Rev. Mol. Cell Biol. 6, 9–20 (2005). 10.1038/nrm1547

49 Fischer, Eric S. et al./person-group>. The Molecular Basis of CRL4DDB2/CSA Ubiquitin Ligase Architecture, Targeting, and Activation. Cell 147, 1024–1039 (2011). 10.1016/j.cell.2011.10.035

50 Zou, J. et al./person-group>. The novel protein homeostatic modulator BTX306 is active in myeloma and overcomes bortezomib and lenalidomide resistance. J. Mol. Med. 98, 1161–1173 (2020). 10.1007/s00109-020-01943-6

51 Nishiguchi, G. et al./person-group>. Identification of Potent, Selective, and Orally Bioavailable Small-Molecule GSPT1/2 Degraders from a Focused Library of Cereblon Modulators. J. Med. Chem. 64, 7296–7311 (2021). 10.1021/acs.jmedchem.0c01313

52 Powell, C. E. et al./person-group>. Selective Degradation of GSPT1 by Cereblon Modulators Identified via a Focused Combinatorial Library. ACS Chem. Biol. 15, 2722–2730 (2020). 10.1021/acschembio.0c00520

53 Teng, M. et al./person-group>. Development of PDE6D and CK1α Degraders through Chemical Derivatization of FPFT-2216. J. Med. Chem. 65, 747–756 (2022). 10.1021/acs.jmedchem.1c01832

54 Park, S.-M. et al./person-group>. Dual IKZF2 and CK1α degrader targets acute myeloid leukemia cells. Cancer Cell 41, 726–739.e711 (2023). 10.1016/j.ccell.2023.02.010

55 Huang, L. et al./person-group>. Development of Oral, Potent, and Selective CK1α Degraders for AML Therapy. JACS Au 4, 4423–4434 (2024). 10.1021/jacsau.4c00762

56 Singhal, S. et al./person-group>. Antitumor Activity of Thalidomide in Refractory Multiple Myeloma. N. Engl. J. Med. 341, 1565–1571 (1999). doi:10.1056/NEJM199911183412102

