## Supplemental information for "A Cryptic Interfacial Pocket Uncovered in Full CRL4^CRBN^–IKZF3 Ubiquitylation Complex Enhances IMiD Efficacy"

#### Materials and Methods

##### Cloning and plasmid construction

The cDNA of human CUL4A (Uniprot entry Q13619, full length, amino acid residues 1–759), RBX1 (Uniprot entry P62877, full length, amino acid residues 1–108), Nedd8 (Uniprot entry Q15843, full length, amino acid residues 1–81), UbcH5a (Uniprot entry P51668, full length, amino acid residues 1–147), and IKZF3 (Uniprot entry Q9UKT9, tandem ZF2–ZF3, amino acid residues 142–198) were synthesized with sequence optimization for *Escherichia coli* recombinant expression by GenScript Biotech (Nanjing, China). The cDNA of human DDB1 (Uniprot entry Q16531, full length, amino acid residues 1–1,140), CRBN (Uniprot entry Q96SW2, amino acid residues 40–442) were synthesized with sequence optimization for HEK293F overexpression by GenScript Biotech (Nanjing, China).

RBX1 and CUL4A were cloned into our home-modified pET29a-YS14 polycistronic vector (*1*) with an N-terminal $His_6$–HRV3C-tag and an N-terminal MysB–HRV3C–StrepII-tag, respectively, according to a reported strategy (*2*). Nedd8 was cloned into a pET22b vector without any tag. UbcH5a or IKZF3 tandem ZF2–ZF3 was individually cloned into a pET28a vector with an N-terminal $His_6$–HRV3C-tag or N-terminal MBP–HRV3C–HA-tag, respectively. DDB1 or CRBN was individually cloned into our home-modified pGX033 vector (pcDNA3.1 backbone with EF1a promoter and WPRE element) (*1*) with an N-terminal FLAG-tag or $His_8$–HRV3C-tag, respectively. The Nedd8 E1 enzyme complex NAE1/UBA3 was cloned into a pGEX-6P-1 vector with an N-terminal GST-tag on NAE1 for polycistronic expression. The Nedd8 E2 enzyme UBE2M was cloned into a pGEX-6P-1 vector with an N-terminal GST-tag. Mutations or truncations were introduced by standard site-directed PCR mutagenesis or homologous recombination.

##### Protein expression and purification

The RBX1/CUL4A complex was polycistronically expressed and purified through a two-step affinity chromatography process. Plasmids containing RBX1/CUL4A were transformed into *Escherichia coli* BL21(DE3) cells (Transgene). Cells were cultured at 37 °C in Luria–Bertani (LB) medium supplemented with 50 μg $mL^{-1}$ kanamycin until reaching an optical density at 600 nm ($OD_{600}$) of 0.8, then induced overnight at 16 °C with 0.2 μM $ZnCl_2$ and 0.5 mM isopropyl β-D-thiogalactopyranoside (IPTG). After centrifugation, cells were lysed by sonication in ice-cold CRL lysis buffer containing 50 mM Tris-HCl (pH 7.5), 300 mM NaCl, 20 mM imidazole, 10% (v/v) glycerol, and 1 mM phenylmethylsulfonyl fluoride (PMSF). Cell lysates were clarified by centrifugation at 12,000 rpm for 30 min at 4 °C, and the supernatant was subsequently loaded onto a Ni-NTA affinity chromatography column. The $His_6$–HRV3C–RBX1/MysB–HRV3C–StrepII–CUL4A protein complex was eluted using Ni elution buffer (50 mM Tris-HCl, pH 7.5, 300 mM NaCl, 300 mM imidazole, and 10% (v/v) glycerol). The eluted proteins were then dialyzed overnight at 4 °C against a buffer containing 50 mM Tris-HCl (pH 7.5), 50 mM NaCl, 5% (v/v) glycerol, and 1 mM dithiothreitol (DTT), during which the $His_6$-tags on RBX1 and MysB-tags on CUL4A were removed by the addition of HRV3C protease. Dialysed proteins were incubated with streptactin resins (Smart-Lifesciences) for 2 h at 4 °C. After extensive wash, RBX1/StrepII–CUL4A complexes were eluted with buffer containing 50 mM Tris-HCl (pH 7.5), 50 mM NaCl, 5% (v/v) glycerol, 1 mM DTT, and 2.5 mM D-desthiobiotin (IBA Lifesciences). The RBX1/StrepII-CUL4A complex was further purified using a Superdex 200 10/300 GL size-exclusion column (GE Healthcare) pre-equilibrated in CRL SEC buffer (20 mM HEPES, pH 7.5, 150 mM NaCl, 2% (v/v) glycerol, 1 mM DTT).

The DDB1/CRBN complex and its mutants were expressed in mammalian cells. HEK293F cells were cultured in Union-293 chemically defined medium (Union-Biotech) to a density of approximately $1.5\times10^6$ cells $mL^{-1}$. DDB1 and CRBN plasmids were transiently co-transfected using polyethyleneimine (PEI, Polysciences). A total of 1.5 mg plasmids and 4.5 mg PEI were mixed in 50 mL of fresh medium for 15–20 min and then added to 1 L of cell culture. After 60 hours of incubation at 37 °C in a shaker with 5% $CO_2$, the transfected cells were collected by centrifugation. The resulting cell pellets were lysed by sonication following resuspension in ice-cold 293F lysis buffer (25 mM HEPES, pH 8.0, 200 mM KCl, 10% (v/v) glycerol, 0.01% (v/v) Triton X-100) supplemented with EDTA-free protease inhibitor tablets (Roche, cOmplete tablets). Cellular debris was cleared by centrifugation at 20,000 × g for 45 min at 4 °C, and the resulting supernatant was incubated with anti-FLAG affinity resin for 2 h at 4 °C. The bound DDB1/CRBN complex was eluted using 293F lysis buffer supplemented with 0.75 mg $mL^{-1}$ FLAG peptide. The DDB1/CRBN complex was subsequently purified by size-exclusion chromatography using either a Superdex 200 10/300 GL size-exclusion column (GE Healthcare) pre-equilibrated with buffer containing 25 mM HEPES (pH 7.5), 200 mM KCl, and 1 mM DTT.

Nedd8 was expressed in *Escherichia coli* Rosetta(DE3) cells (Solarbio). The cells were grown in LB medium supplemented with 50 μg $mL^{-1}$ ampicillin at 37 °C until reaching an $OD_{600}$ of 1.0, and protein expression was induced by adding 0.6 mM IPTG, followed by overnight incubation at 18 °C. Cells were harvested by centrifugation and lysed via sonication in N8 lysis buffer (50 mM HEPES, pH 7.5, 150 mM NaCl). The lysate was subjected to ultracentrifugation at 12,000 rpm for 30 min at 4 °C. The resulting pellet was resuspended in unfolding buffer (20 mM Tris-HCl, pH 7.8, 6 M guanidine hydrochloride (Gn-HCl), 1 mM DTT), and a second round of ultracentrifugation was performed. The supernatant was dialyzed overnight against double-distilled water ($ddH_2O$) containing 0.1% (v/v) trifluoroacetic acid (TFA). After dialysis, the mixture was clarified again by ultracentrifugation, and the resulting supernatant was purified by semi-preparative reverse-phase high-performance liquid chromatography (RP-HPLC), followed by electrospray ionization mass spectrometry (ESI-MS) characterization. The purified Nedd8 protein was then lyophilized into powder. For refolding, the lyophilized Nedd8 protein was first dissolved in unfolding buffer and then gradually diluted by adding N8 lysis buffer. Final purification was carried out using a Superdex 75 Increase 10/300 GL column (GE Healthcare) pre-equilibrated with N8 lysis buffer.

IKZF3 tandem ZF2–ZF3 motif and its mutants were expressed in *Escherichia coli* BL21(DE3) cells (Transgene). Cell cultures were grown at 37 °C in LB medium supplemented with 50 μg $mL^{-1}$ kanamycin until the $OD_{600}$ reached 1.0. IKZF3 expression was then induced by adding 0.2 μM $ZnCl_2$ and 0.5 mM IPTG, followed by overnight incubation at 16 °C. Cells were harvested by centrifugation and lysed by sonication in ice-cold IKZF lysis buffer composed of 50 mM Tris-HCl (pH 7.5), 200 mM NaCl, 10% (v/v) glycerol, 1 mM DTT, 50 μM $ZnCl_2$, and 1 mM PMSF. After centrifugation, the clarified supernatant was applied to an MBP-Tag Dextrin Resin (PurKine™, Abbkine) for 4 h at 4 ℃. After extensive wash by IKZF wash buffer (50 mM Tris-HCl, pH 7.5, 200 mM NaCl, 10% (v/v) glycerol, 1 mM DTT, and 50 μM $ZnCl_2$), HRV3C protease was introduced into the resin suspension in IKZF wash buffer and incubated under gentle agitation overnight at 4 ℃ to cleave IKZF3 ZF2–ZF3 proteins from the resin. Cleaved IKZF3 ZF2–ZF3 proteins were further purified using a Superdex 75 Increase 10/300 GL column (GE Healthcare) pre-equilibrated with CRL SEC buffer.

UbcH5a was expressed in *Escherichia coli* BL21(DE3) cells (Transgene) and purified by Ni affinity chromatography, HRV3C protease cleavage, and size-exclusion chromatography (Superdex 75 10/300 GL column, GE Healthcare).

NAE1/UBA3 and UBE2M were individually expressed and purified from *Escherichia coli* BL21(DE3) cells (Transgene). For NAE1/UBA3, the cells were cultured in LB medium supplemented with 50 μg $mL^{-1}$ ampicillin at 37 °C to an $OD_{600}$ of 1.0 and induced by the addition of 0.4 mM IPTG at 16 °C overnight. Cells were harvested by centrifugation and lysed by sonication in GST lysis buffer (25 mM HEPES, pH 7.5, 150 mM NaCl, 10% (v/v) glycerol, 1 mM DTT). Lysates were ultracentrifuged at 12,000 rpm for 30 min at 4 °C, and the resulting supernatant was loaded onto a Glutathione Beads 4FF (Lableads). The NAE1/UBA3 proteins were eluted using GST lysis buffer supplemented with 30 mM reduced glutathione (GSH), followed by HRV3C protease cleavage overnight. The protein was further purified by ion exchange chromatography using a 5 mL HiTrap Q column. The UBE2M protein was expressed and purified following the same procedure as NAE1/UBA3, with the exception that the final purification step was performed using a Hiload 16/600 SD75 pg column (GE Healthcare) pre-equilibrated with a buffer containing 50 mM HEPES (pH 7.5 and 150 mM NaCl.

Ubiquitin (Ub) and its mutants were expressed and purified as previously described(*3*). Briefly, proteins were overexpressed in *Escherichia coli* BL21(DE3) cells (Transgene). Following cell harvest, pellets were resuspended in $ddH_2O$ and lysed by sonication. The lysates were treated with 1% (v/v) perchloric acid to precipitate contaminants and subsequently clarified by centrifugation. The resulting supernatant was dialysed against $ddH_2O$ containing 0.1% (v/v) TFA overnight and then purified using a Source15S cation exchange column (GE Healthcare). Fractions corresponding to Ub were pooled and dialyzed against a buffer containing 20 mM HEPES (pH 7.5) and 150 mM NaCl.

**Reconstitution of $_{N8}$CRL4$^{CRBN}$ E3 ligase complex**

For neddylation of RBX1/CUL4A, 0.8 μM NAE1/UBA3, 6.5 μM UBE2M, 20 μM Nedd8, and 2.5 μM RBX1/CUL4A were mixed in a buffer containing 50 mM Tris-HCl (pH 7.5), 150 mM NaCl, 1 mM DTT, 5 mM adenosine 5'-phosphate (ATP), and 5 mM $MgCl_2$. The reaction mixture was incubated at 27 °C for 4 h. Following confirmation of complete neddylation of CUL4A by SDS–PAGE analysis, the DDB1/CRBN complex was subsequently added to the reaction mixture at a molar ratio corresponding to 0.75 equivalents relative to RBX1/CUL4A. The mixture was further incubated at 4 °C for 2 h and purified by a Superdex 200 10/300 GL column (GE Healthcare) pre-equilibrated with CRL SEC buffer. Fractions containing the $_{N8}$CRL4$^{CRBN}$ E3 ligase complex were pooled, analyzed by SDS–PAGE, concentrated to ~2.6 mg $mL^{-1}$, and stored at –80 °C.

**Fluorescent labeling of ubiquitin**

Ub-MCQ (a ubiquitin variant with an additional cysteine inserted following the N-terminal methionine) was concentrated to approximately 20 mg $mL^{-1}$ in buffer containing 20 mM HEPES (pH 7.5) and 150 mM NaCl. Oregon Green™ 488 (OG488) maleimide (1.2 equivalents, prepared as a 20 mg $mL^{-1}$ stock in dimethyl sulfoxide (DMSO), purchased from Thermo Fisher Scientific) was then added, and the pH of the solution was adjusted to 7.4. The reaction was carried out at 4 °C for 24 h and subsequently quenched by the addition of DTT to a final concentration of 100 mM. The fluorescently labeled ubiquitin (Ub$^{OG488}$) was purified via size-exclusion chromatography using a Superdex 75 column (GE Healthcare) pre-equilibrated with a buffer containing 20 mM HEPES (pH 7.5), 150 mM NaCl, 2% (v/v) glycerol, and 1 mM DTT.

**Chemical synthesis of IKZF3–Ub ABP**

The IKZF3–Ub ABP was synthesized following previously reported protocols (*4, 5*) with some modifications (fig. S2B). Initially, lyophilized Ub(1–75)-MesNa (3 mg $mL^{-1}$, prepared following the described protocol (*6*)) was dissolved in reaction buffer (20 mM HEPES, pH 7.5, 6 M Gn-HCl). Subsequently, 400 mM (E)-3-[2-(bromomethyl)1,3-dioxolan-2-yl]prop-2-en-1-amine (BmDPA, Bide Pharmatech Co., Ltd) was added to the solution without adjusting the pH. The reaction mixture was incubated in the dark at 37 °C for 2 h, followed by purification via semi-preparative RP-HPLC and characterization by ESI-MS. The purified product was lyophilized and dissolved at a concentration of 0.5 mg $mL^{-1}$ in deprotection buffer (20 mM HEPES, 6 M Gn-HCl, 54% (v/v) TFA, 40 mM p-toluenesulfonic acid). Deprotection was performed in the dark at 37 °C for 1 h. The resulting Ub–BmDPA was then purified by semi-preparative RP-HPLC, confirmed by ESI-MS analysis (fig. S2C), and lyophilized into powder. Refolded Ub–BmDPA was concentrated to ~10 mg $mL^{-1}$.

The IKZF3 ZF2–ZF3 sequence contains a non-zinc-coordinating cysteine residue at position 183 (C183). To ensure site-specific reactivity at the K166C position, C183 was mutated to threonine (C183T). Purified IKZF3 ZF2–ZF3 C183T/K166C proteins were mixed with Ub–BmDPA to initiate the nucleophilic substitution reaction. The reaction mixture was separated on a Superdex 75 Increase column (GE Healthcare) pre-equilibrated with a buffer containing 20 mM HEPES (pH 7.5), 150 mM NaCl, 2% (v/v) glycerol. Fractions containing IKZF3–Ub ABP were pooled, analyzed by SDS–PAGE (fig. S2D), concentrated to 0.5 mg $mL^{-1}$, and stored at –80 °C.

**IMiD chemicals**

Eight IKZF1/3-targeting IMiD molecules used in this study were purchased from commercial suppliers and used without further purification. Pomalidomide (CC-4047, Cat# HY-111109) and lenalidomide (CC-5013, Cat# HY-A0003) were purchased from MedChemExpress. Thalidomide (Cat# S1193), avadomide (CC-122, Cat# S7892), iberdomide (CC-220, Cat# S8760), mezigdomide (CC-92480, Cat# S8975), golcadomide (CC-99282, Cat# E1212), and cemsidomide (CFT7455, Cat# E1184) were purchased from Selleck. The IMiD chemicals were supplied as dry powders and individually dissolved in anhydrous DMSO to prepare 10 mM stock solutions. Aliquots were stored at –80 °C for future use.

***In vitro* ubiquitylation assays**

Generally, assays were conducted in a reaction mixture containing human UBA1, UbcH5a, $_{N8}$CRL4$^{CRBN}$ E3 ligase complex, IKZF3 tandem ZF2–ZF3 motifs, Ub$^{OG488}$, and IMiDs in Ub assay buffer (50 mM Tris-HCl, pH 7.5, 150 mM NaCl, 10 μM $ZnCl_2$, 2 mM $MgCl_2$, 3 mM ATP, and 1 mM DTT). In all experiments, the final concentration of DMSO introduced by the IMiD stock solution was maintained at 2% (v/v) in the reaction mixture. Before use, all enzymes — including relevant fragments and mutants — were analyzed by SDS–PAGE (SurePAGE gels, GenScript Biotech). Reactions were terminated by the addition of 4 × LDS loading buffer (Thermo Fisher Scientific) supplemented with 100 mM DTT, followed by heating at 95 °C for 5 min. Reactions were then resolved by SDS–PAGE (NuPAGE gels, Thermo Fisher Scientific).

UBA1, UbcH5a, $_{N8}$CRL4$^{CRBN}$ E3 ligase complex, IMiD, and IKZF3 ZF2–ZF3 were first mixed in Ub assay buffer and incubated on ice for 5 min. Then Ub$^{OG488}$ was introduced into the mixture to initiate ubiquitylation reactions.

For the IKZF3 and CRBN mutagenesis assays shown in Figs. 3J and 3K, 1.0 μM UBA1, 2.0 μM UbcH5a, 0.10 μM N8CRL4$^{CRBN}$ E3 ligase complex or its CRBN H103A mutant, 5.0 μM IKZF3 ZF2–ZF3 or its R186A mutant, 10.0 μM Ub$^{OG488}$, and 2.0 μM IMiD were used. Reactions were incubated at 37 °C and quenched at $t$ = 10 min. Samples were resolved on 12% NuPAGE Bis-Tris gels (Thermo Fisher Scientific).

Experiments were performed in technical triplicate ($n$ = 3).

#### *In vivo* degradation assays

HEK293T cells were maintained in Dulbecco's modified Eagle's medium (DMEM, Thermo Fisher Scientific) supplemented with 10% (v/v) fetal bovine serum (FBS, Thermo Fisher Scientific) and 0.01% (v/v) penicillin-streptomycin stock (10,000 U mL$^{-1}$, Thermo Fisher Scientific). HEK293T cells plated in 6-well plates were transfected with pcDNA3.1 plasmids (2.5 μg per well) containing N-terminal FLAG-tagged full-length IKZF3 (IKZF3$^{FL}$) or zinc finger deletion constructs (IKZF3$^{\Delta ZF2}$ or IKZF3$^{\Delta ZF3}$) using Lipofectamine 3000 reagent (Thermo Fischer Scientific). After 24 h of transfection, the culture medium was replaced with fresh medium supplemented with individual IMiD at a final concentration of 5 μM for each compound. In all wells, the final concentration of DMSO was maintained at 0.005% (v/v) in the medium. After 48 h of IMiD treatment, cells were washed with ice-cold phosphate buffered saline (PBS, Thermo Fischer Scientific) and harvested using cell scrapers. Harvested cells were pelleted by centrifugation. Cells were processed using Minute Total Protein Extraction Kit for Animal Cultured Cells and Tissues (Cat# SD-001/SN-002, Invent Biotechnologies, Inc.) following the protocol provided by the supplier to prepare the cell lysates. Total proteins in cell lysates were immediately measured using Pierce BCA Protein Assay Kits (Thermo Fischer Scientific) and analyzed by western blots as previously described (*1*). Briefly, following the membrane transfer procedure, the polyvinylidene difluoride (PVDF) membrane (Immobilon-P, Merck Millipore) was horizontally cut at the 50 kDa molecular weight marker. After blocking, the upper portion (> 50 kDa) was probed with a rabbit monoclonal anti-FLAG antibody (Cell Signaling Technology, Cat# D6W5B, 1:1,000 diluted), while the lower portion (< 50 kDa) was probed with a rabbit monoclonal anti-β-actin antibody (Cell Signaling Technology, Cat# 4970S, 1:4,000 diluted). An HRP-conjugated goat-anti-rabbit IgG antibody (2031, Huaxingbio) was used as the secondary antibody. The membranes were visualized using the Clarity Western ECL substrate (Bio-Rad) and imaged with a ChemiDoc MP Imaging System (Bio-Rad). Experiments were performed in technical duplicate ($n$ = 2).

#### Cryo-EM sample preparation

UbcH5a was added to the reaction mixture that contained N8CRL4$^{CRBN}$ E3 ligase complex, IKZF3–Ub ABP, and IMiD reagent. In all reaction mixtures, the final concentration of DMSO was maintained under 0.3% (v/v). The trapped ubiquitylation assembly complexes were purified using a Superdex 200 Increase 5/150 GL column or a Superose 6 Increase 5/150 GL column (GE Healthcare) pre-equilibrated with CRL REC buffer (20 mM HEPES, pH 7.5, 75 mM NaCl, 1 mM TCEP). Peak fractions were pooled, analyzed by SDS–PAGE, and immediately subjected to cryo-EM sample preparation.

A 3.5 μL aliquot of the sample was applied to glow-discharged holey gold grids (Quantifoil R1.2/1.3, Au 300 mesh) and incubated at 8 °C under 100% humidity for 60 seconds. The initial sample was removed from the grid using a pipette, followed by the application of a new 3.5 μL aliquot for an additional 60-second incubation. Grids were blotted and rapidly plunge-frozen in liquid ethane using a Vitrobot (Thermo Fisher Scientific).

**Cryo-EM data collection**

A total of 13,668 cryo-EM micrographs of the pomalidomide dataset were collected on a 300 kV Titan Krios microscope configured with a Gatan K3 direct electron detector camera (Tsinghua University, Branch of China National Center for Protein Sciences, Beijing) using the AutoEMation software (*7*). Micrographs of the pomalidomide dataset were recorded at a pixel size of 1.10 Å with a magnification of 81,000× and a dose of 50 $e^-$ $Å^{-2}$ for 32 frames with a defocus range of –1.0 to –1.8 µm. A total of 7,706 (the thalidomide dataset), 7,621 (the lenalidomide dataset), 8,474 (the avadomide dataset), 8,097 (the iberdomide dataset), 25,882 (the mezigdomide dataset), 8,567 (the golcadomide dataset), and 7,205 (the cemsidomide dataset) cryo-EM micrographs were collected on a 300 kV Titan Krios G4 microscope configured with a Falcon4 direct electron detector camera (Shuimu BioSciences, Hangzhou) using the EPU software (Thermo Fisher Scientific). For the mezigdomide dataset, micrographs were collected at a magnification of 96,000× with a pixel size of 0.830 Å. A total electron dose of 50 $e^-$ $Å^{-2}$ was applied over 32 frames, with a defocus range of –1.2 to –2.0 µm. Micrographs for the remaining six datasets (the thalidomide, lenalidomide, avadomide, iberdomide, golcadomide, and cemsidomide datasets) were acquired at 75,000× magnification, corresponding to a pixel size of 1.059 Å, under identical exposure conditions.

**Cryo-EM data processing**

All datasets were processed using RELION v3.1.1 (*8*). Motion correction, contrast transfer function (CTF) estimation, manual or automated particle picking, and particle extraction were performed. Subsequent steps included iterative 2D and 3D classifications, mask generation, 3D auto-refinement, and postprocessing. Data processing flow charts are detailed in fig. S3 (the mezigdomide dataset) and Data S1 (the thalidomide, lenalidomide, pomalidomide, avadomide, iberdomide, golcadomide, and cemsidomide datasets). To improve the cryo-EM density of IKZF ZF3, a mask and focused 3D classifications were applied to the CRBN TBD–Lon–IKZF3–UbcH5a–RBX1 region. Particles exhibiting the best ZF3 features were selected and used for the final reconstruction. Cryo-EM data processing statistics are summarized in Tables S1 and S2. Overall resolutions were determined based on the gold standard Fourier shell correlation (FSC) criterion at 0.143, and local resolution estimations were carried out using ResMap v1.1.4.

**Model building, refinement, and validation**

An initial model containing mezigdomide was generated using previously reported models of subunits or subcomplexes: the cullin scaffold containing CUL4A and Nedd8 and the catalytic module containing RBX1, UbcH5a, and Ub (PDB 8B3G) (*9*), the ternary module containing DDB1, CRBN, IKZF3, and mezigdomide (PDB 8D7Z) (*10*). The IKZF1 ZF2 motif in PDB entry 8D7Z was replaced with the AlphaFold2 (*11*) model of the IKZF3 ZF2 motif (Q9UKT9-F1-v4), and the IKZF3 ZF3 motif was further added based on the AlphaFold2 model. These models were manually fit into the mezigdomide cryo-EM map by rigid-body docking using UCSF ChimeraX v1.9 (*12*). The initial model was generated based on the coordinates manually docked in UCSF ChimeraX using COOT v0.8.2 (*13*). Next, one round of real-space refinement using Phenix v1.21.1 (*14*) was performed to yield an initial refined model. The mezigdomide molecule in the initially refined model was individually replaced with each of the other seven IMiDs. The restraints and coordinates of five IMiDs were derived from previously reported crystal structures of binary or ternary complexes: thalidomide (PDB 4CI1) (*15*), lenalidomide (PDB 4CI2) (*15*), pomalidomide (PDB 4CI3) (*15*), avadomide (PDB 7PSO) (*16*), and iberdomide (PDB 8D80) (*10*). As no binary/ternary structures containing golcadomide or cemsidomide had been reported when we conducted model building, restraints and initial PDB models for golcadomide and cemsidomide were generated using the eLBOW program in Phenix-1.21.1 (*14*). Iterative cycles of real-space refinement using Phenix v1.21.1 (*14*) and manual model adjustment with COOT v0.8.2 (*13*) were performed until satisfactory map-to-model correlation and geometry were obtained. Protein backbone segments and side chains not supported by cryo-EM density were manually removed. Tables S1 and S2 provide a summary of all statistics related to model building, refinement, and validations. All structural interpretations, visualizations were performed using UCSF ChimeraX v1.9 (*12*).

***In silico* docking**

Molecular docking was performed following standard protocols using Maestro 14.1 (Schrödinger, New York). The mezigdomide-organized full ubiquitylation complex resolved in this study and previously reported IKZF1–mezigdomide–CRBN/DDB1 (PDB 8D7Z) ternary complex were prepared using the Protein Preparation programme. The receptor grid was centered on mezigdomide for grid generation. The LigPrep programme was employed to generate conformers of TMX-4116, dCK1α-2, SJ3149, DEG-35, eragidomide, SJ6986, ZHX-1-161, BTX306, CC-885 and mezigdomide analog molecules. Docking was carried out using the Glide module in standard precision mode. Docking results were evaluated based on the docking score and visualized in UCSF ChimeraX v1.9 (*12*).

**Protein bands quantification and data analysis**

For *in vitro* ubiquitylation assays, protein band quantification was performed as previously described (*1*). Gels were imaged using a ChemiDoc MP Imaging System (Bio-Rad). Initially, the Fluorescein setting was used to visualize fluorescent proteins, followed by the Coomassie Far Red Epi mode to detect protein ladders. The same gel was subsequently stained with Coomassie Brilliant Blue (CBB). Fluorescent and Coomassie Far Red Epi images were merged to estimate the molecular weights of the fluorescent proteins. Quantitative analysis of protein bands in both unmerged fluorescent gel images and CBB-stained gel images was carried out using Image Lab 6.0.1 (Bio-Rad). Boxes were drawn around the ubiquitylated and unmodified IKZF3 bands in the CBB-stained gel images, followed by background signal subtraction. Intensities of protein bands in each lane were calculated. Fractions of total IKZF3 (%) were calculated by dividing the band intensity of the ubiquitylated IKZF3 species (e.g., IKZF3–Ub) by the total band intensity of all IKZF3 species in the same lane.

For *in vivo* degradation assays, boxes were placed around the IKZF3 bands and actin bands in the blot images, and background signals were subtracted. IKZF3 remaining (%) was calculated by dividing the band intensity of the IKZF3 band by the intensity actin band in the same lane.

### Figure S1

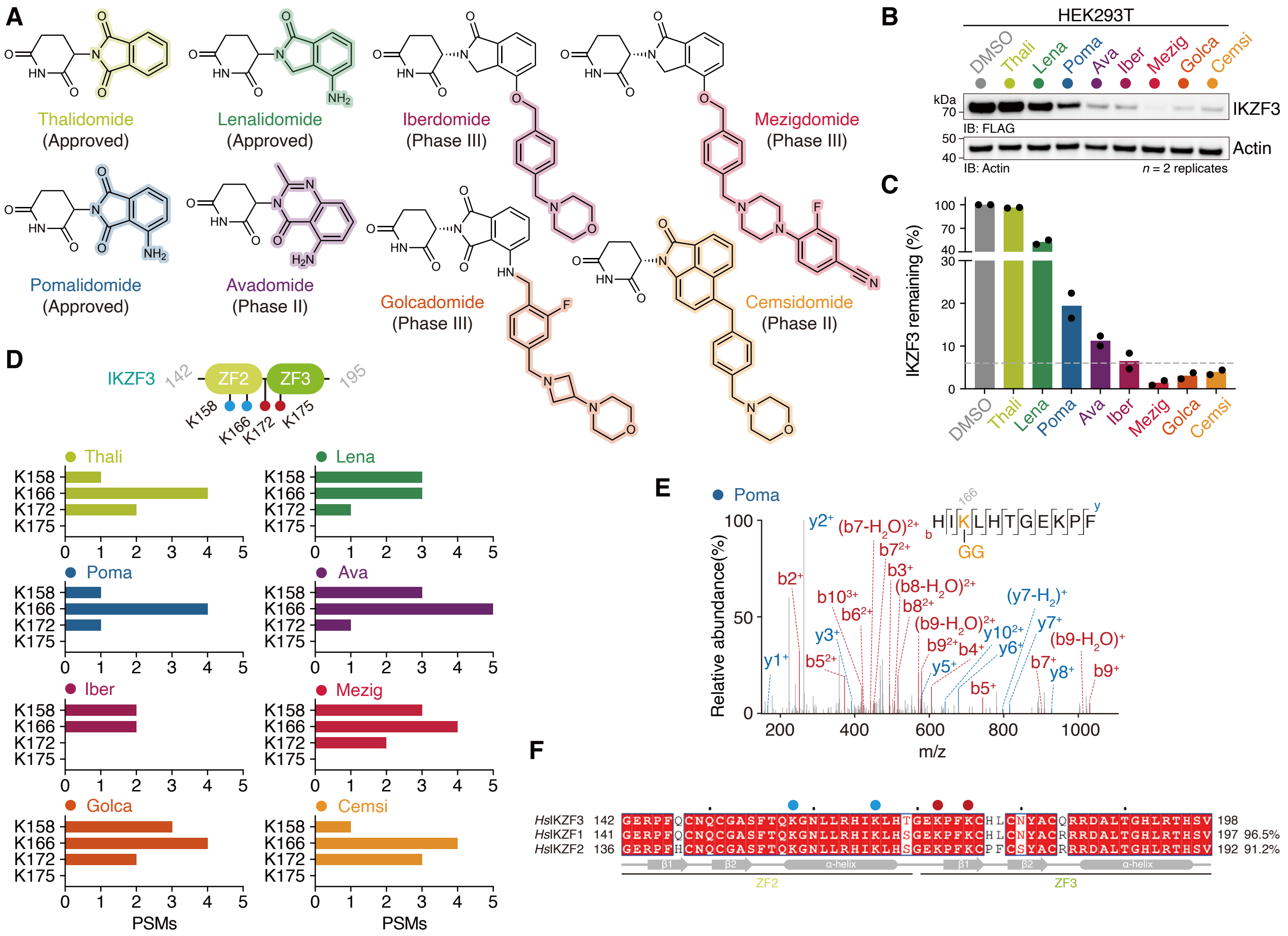
A
Thalidomide
(Approved)
Lenalidomide
(Approved)
Iberdomide
(Phase III)
Mezigdomide
(Phase III)
Pomalidomide
(Approved)
Avadomide
(Phase II)
Golcadomide
(Phase III)
Cemsidomide
(Phase II)
B
HEK293T
DMSO Thali Lena Poma Ava Iber Mezig Golca Cemsi
kDa
70
IKZF3
IB: FLAG
50
40
Actin
IB: Actin
n = 2 replicates
C
IKZF3 remaining (%)
DMSO Thali Lena Poma Ava Iber Mezig Golca Cemsi
D
IKZF3 142 ZF2 ZF3 195
K158 K166 K172 K175
Thali
Lena
Poma
Ava
Iber
Mezig
Golca
Cemsi
PSMs
E
Poma
HIKLHTGEKPF
GG
Relative abundance(%)
m/z
F
HsIKZF3 142 GERPFQCNQCGASFTQKGNLLRHIKLHTGEKPFKCHLCNYACQRRDALTGHLRTHSV 198
HsIKZF1 141 GERPFQCNQCGASFTQKGNLLRHIKLHSGEKPFKCHLCNYACRRRDALTGHLRTHSV 197 96.5%
HsIKZF2 136 GERPFHCNQCGASFTQKGNLLRHIKLHSGEKPFKCPFCSYACRRRDALTGHLRTHSV 192 91.2%
β1 β2 α-helix β1 β2 α-helix
ZF2
ZF3

**Fig. S1. Approved and clinical IMiDs exhibit distinct degradation efficiencies and site-selective ubiquitylation of IKZF3.**

(**A**) Chemical structures of eight approved and clinical IKZF3-targeting IMiDs, with differences in each molecule highlighted by color.

(**B**) A representative western blot image of *in vivo* IKZF3 degradation assay in the presence of each IMiD (1 μM final concentration).

(**C**) Bar graph showing quantified IKZF3 degradation data from $n = 2$ technical replicates.

(**D**) Bar graphs showing the Peptide-Spectrum Matches (PSMs) of peptides containing lysine residues (K158, K166, K172, or K175) modified with GG remnant.

(**E**) A representative tandem mass spectrum (MS/MS spectrum) for a peptide containing a GG remnant modified on IKZF3 K166 derived from *in vitro* ubiquitylation assays using pomalidomide.

(**F**) Sequence alignment of tandem ZF2–ZF3 motifs in human Ikaros family transcription factors IKZF3, IKZF1, and IKZF2. Consensus lysines on the ZF2 or ZF3 motif are indicated by blue or red dots, respectively.

### Figure S2

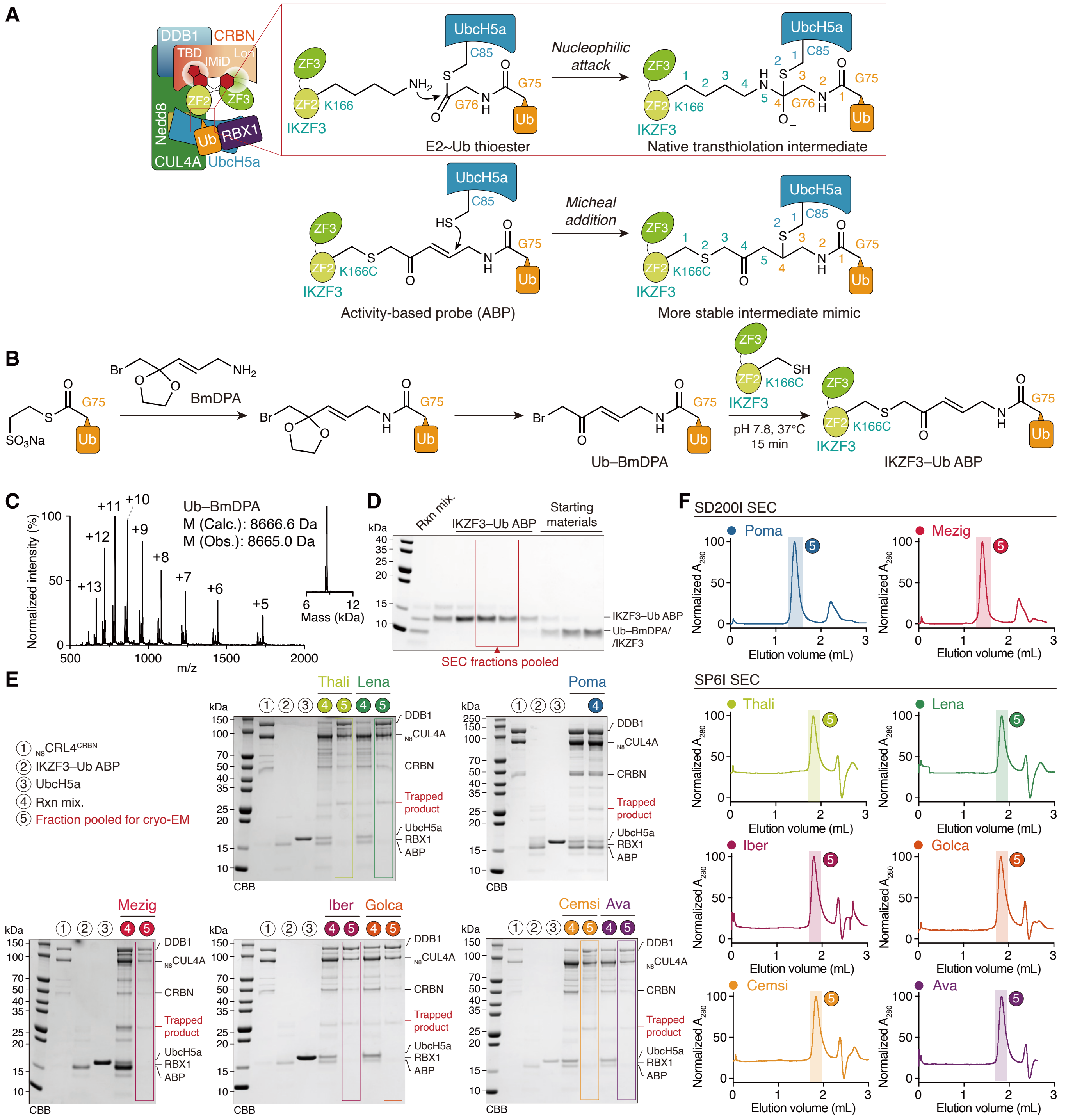
A
DDB1
CRBN
TBD
IMiD
Lon
ZF2
ZF3
Nedd8
Ub
RBX1
CUL4A
UbcH5a
ZF3
ZF2
K166
IKZF3
$NH_2$
UbcH5a
C85
S
O
G76
N
H
O
G75
Ub
E2~Ub thioester
Nucleophilic attack
Native transthiolation intermediate
Activity-based probe (ABP)
HS
K166C
Micheal addition
More stable intermediate mimic
B
$SO_3Na$
BmDPA
Br
$NH_2$
Ub–BmDPA
SH
pH 7.8, 37°C
15 min
IKZF3–Ub ABP
C
Normalized intensity (%)
m/z
+13
+12
+11
+10
+9
+8
+7
+6
+5
Ub–BmDPA
M (Calc.): 8666.6 Da
M (Obs.): 8665.0 Da
Mass (kDa)
D
kDa
Rxn mix.
IKZF3–Ub ABP
Starting materials
IKZF3–Ub ABP
Ub–BmDPA/
/IKZF3
SEC fractions pooled
E
① $_{N8}$CRL4$^{CRBN}$
② IKZF3–Ub ABP
③ UbcH5a
④ Rxn mix.
⑤ Fraction pooled for cryo-EM
Thali
Lena
Poma
Mezig
Iber
Golca
Cemsi
Ava
DDB1
$_{N8}$CUL4A
CRBN
Trapped product
UbcH5a
RBX1
ABP
CBB
F
SD200I SEC
SP6I SEC
Poma
Mezig
Thali
Lena
Iber
Golca
Cemsi
Ava
Normalized $A_{280}$
Elution volume (mL)

**Fig. S2. Implementing activity-based probe to trap the IMiD-induced IKZF3 ubiquitylation assemblies containing fully-activated $_{N8}$CRL4$^{CRBN}$ E3 ligase.**

(**A**) Diagram comparing the formation and geometry of the native transthiolation intermediate for IMiD-induced IKZF3 ubiquitylation and its more stable mimic. The mimic was generated through the Michael addition reaction between the activity-based probe (ABP) and the UbcH5a catalytic cysteine.

(**B**) Synthetic scheme of the IKZF3–Ub ABP. The IKZF3 ZF2–ZF3 K166C was conjugated to the C-terminal G75 of the donor Ub through a BmDPA handle.

(**C**) Electrospray ionization-mass spectrometry (ESI-MS) spectra and deconvoluted MS spectra of Ub–BmDPA with the calculated (Calc.) and observed (Obs.) molecular weight shown.

(**D**) SDS–PAGE analysis of the size-exclusion chromatography (SEC)-purified nucleophilic substitution reaction between Ub–BmDPA and IKZF3 ZF2–ZF3 K166C construct. Pooled fractions containing IKZF3–Ub ABP for structural study are indicated by the red rectangle.

(**E**) SDS–PAGE analysis of chemically-trapped ubiquitylation assembly formation in the presence of eight individual IMiD molecules. Fractions pooled for cryo-EM analysis were indicated by colored rectangles.

(**F**) Size-exclusion chromatograms of ubiquitylation assemblies containing eight individual IMiD molecules. The complex containing pomalidomide or mezigdomide was purified by a Superdex 200 Increase (SD200I) column (Cytiva, GE Healthcare). The complex containing thalidomide, lenalidomide, iberdomide, golcadomide, cemsidomide, or avadomide was purified by a Superose 6 Increase (SP6I) column (Cytiva, GE Healthcare).

### Figure S3

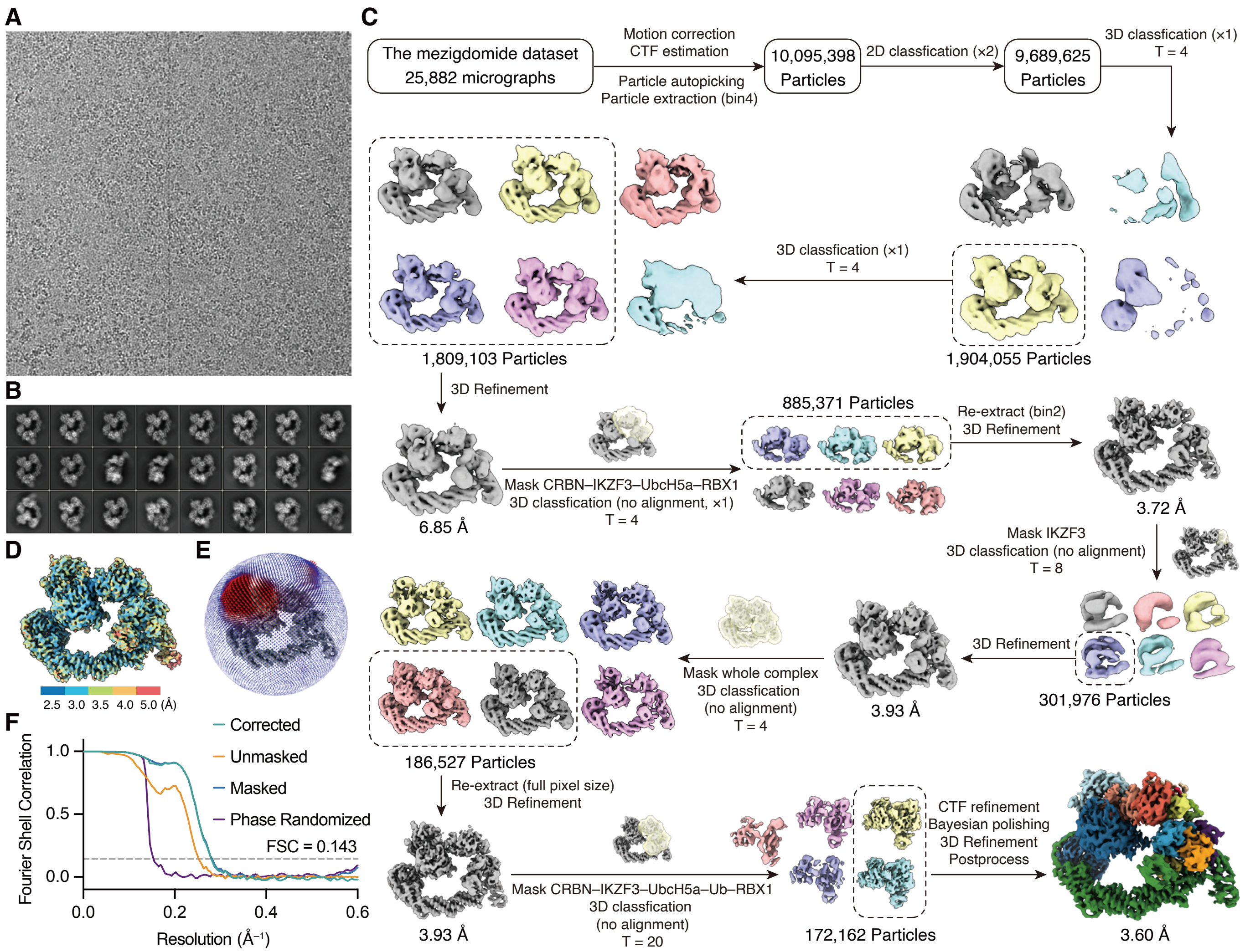

A
B
C
D
E
F
The mezigdomide dataset
25,882 micrographs
Motion correction
CTF estimation
Particle autopicking
Particle extraction (bin4)
10,095,398 Particles
2D classfication (×2)
9,689,625 Particles
3D classfication (×1)
T = 4
3D classfication (×1)
T = 4
1,809,103 Particles
1,904,055 Particles
3D Refinement
6.85 Å
Mask CRBN–IKZF3–UbcH5a–RBX1
3D classfication (no alignment, ×1)
T = 4
885,371 Particles
Re-extract (bin2)
3D Refinement
3.72 Å
Mask IKZF3
3D classfication (no alignment)
T = 8
3D Refinement
301,976 Particles
3.93 Å
Mask whole complex
3D classfication
(no alignment)
T = 4
186,527 Particles
Re-extract (full pixel size)
3D Refinement
3.93 Å
Mask CRBN–IKZF3–UbcH5a–Ub–RBX1
3D classfication
(no alignment)
T = 20
172,162 Particles
CTF refinement
Bayesian polishing
3D Refinement
Postprocess
3.60 Å
2.5 3.0 3.5 4.0 5.0 (Å)
Corrected
Unmasked
Masked
Phase Randomized
FSC = 0.143
Fourier Shell Correlation
1.0
0.5
0.0
0.0
0.2
0.4
0.6
Resolution (Å⁻¹)

**Fig. S3. Cryo-EM data processing flow chart of the mezigdomide-organized ubiquitylation assembly.**

(**A**) A representative micrograph.

(**B**) Representative 2D averages.

(**C**) Data processing workflow.

(**D**) Final cryo-EM map colored by the indicated local resolution.

(**E**) Angular distributions of the final cryo-EM reconstruction.

(**F**) Gold standard Fourier shell correlation (FSC) curves for the final refinement (3.60 Å resolution at FSC = 0.143).

### Figure S4

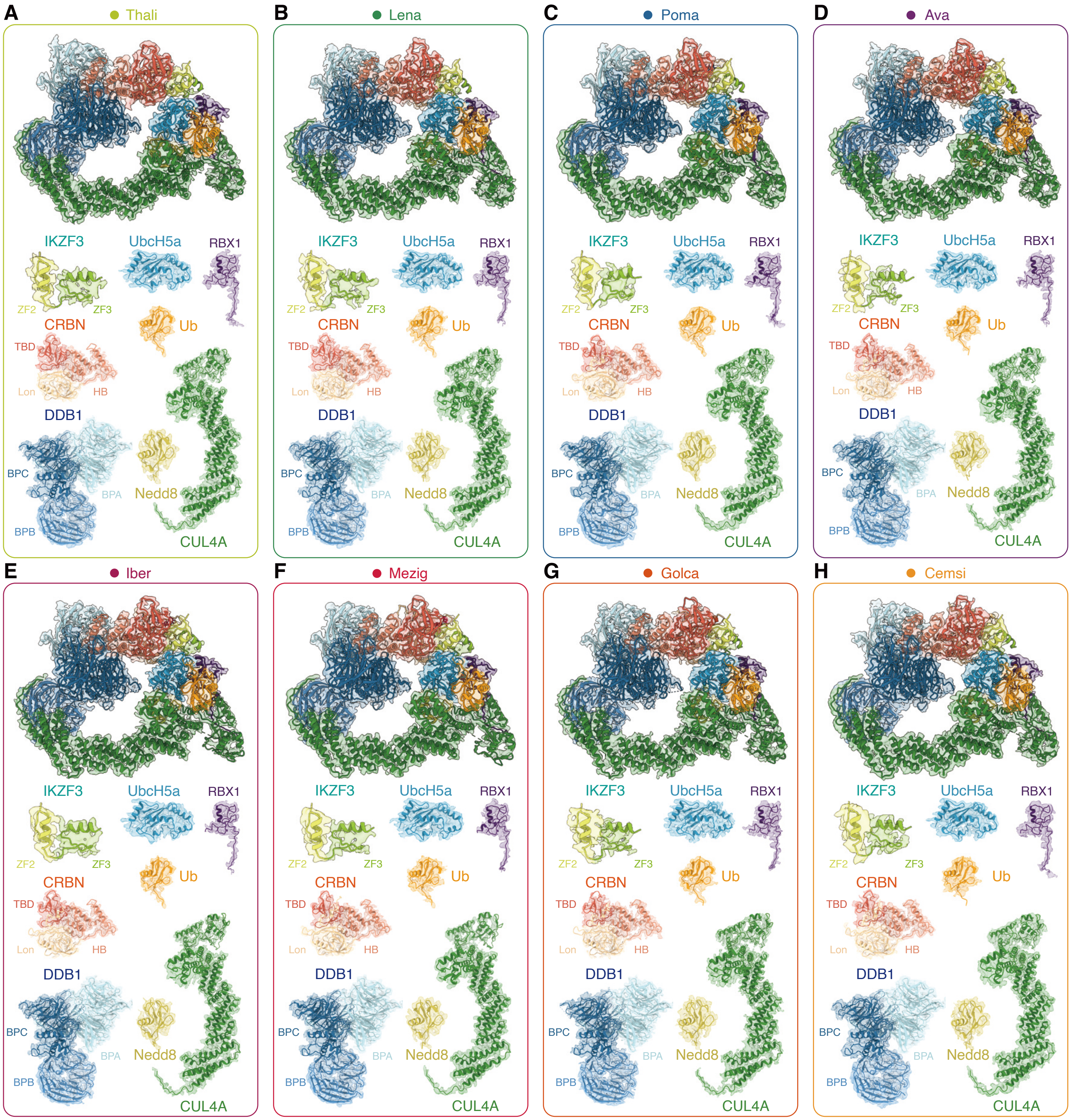
A
Thali
IKZF3
UbcH5a
RBX1
ZF2
ZF3
CRBN
Ub
TBD
Lon
HB
DDB1
BPC
BPA
Nedd8
BPB
CUL4A
B
Lena
C
Poma
D
Ava
E
Iber
F
Mezig
G
Golca
H
Cemsi

**Fig. S4. Cryo-EM density maps and atomic models of various components within the eight IMiD-organized ubiquitylation assemblies.**

(**A**) The thalidomide complex.

(**B**) The lenalidomide complex.

(**C**) The pomalidomide complex.

(**D**) The avadomide complex.

(**E**) The iberdomide complex.

(**F**) The mezigdomide complex.

(**G**) The golcadomide complex.

(**H**) The cemsidomide complex.

### Figure S5

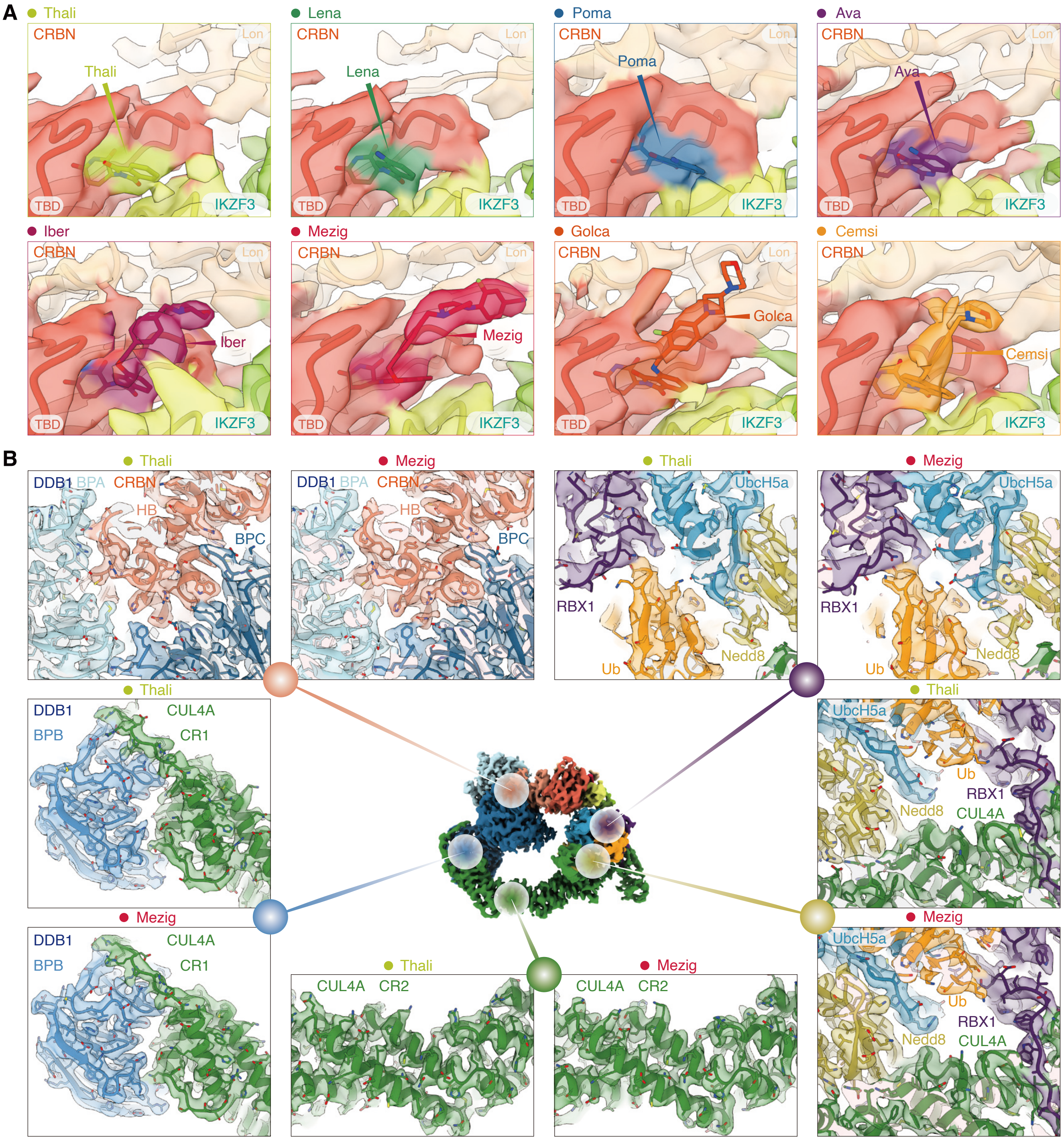
A
Thali
Lena
Poma
Ava
Iber
Mezig
Golca
Cemsi
CRBN
Lon
TBD
IKZF3
B
DDB1
BPA
CRBN
HB
BPC
UbcH5a
RBX1
Ub
Nedd8
CUL4A
BPB
CR1
CR2

**Fig. S5. Cryo-EM density maps and atomic models of key interacting regions of the eight IMiDs-organized ubiquitylation assemblies.**

(**A**) Substrate binding region in which eight individual IMiD molecules bind to the CRBN TBD pocket and IKZF3 ZF2 degron.

(**B**) Representative interfaces within the ubiquitylation assembly. The structures of complexes containing thalidomide or mezigdomide are exhibited as examples.

### Figure S6

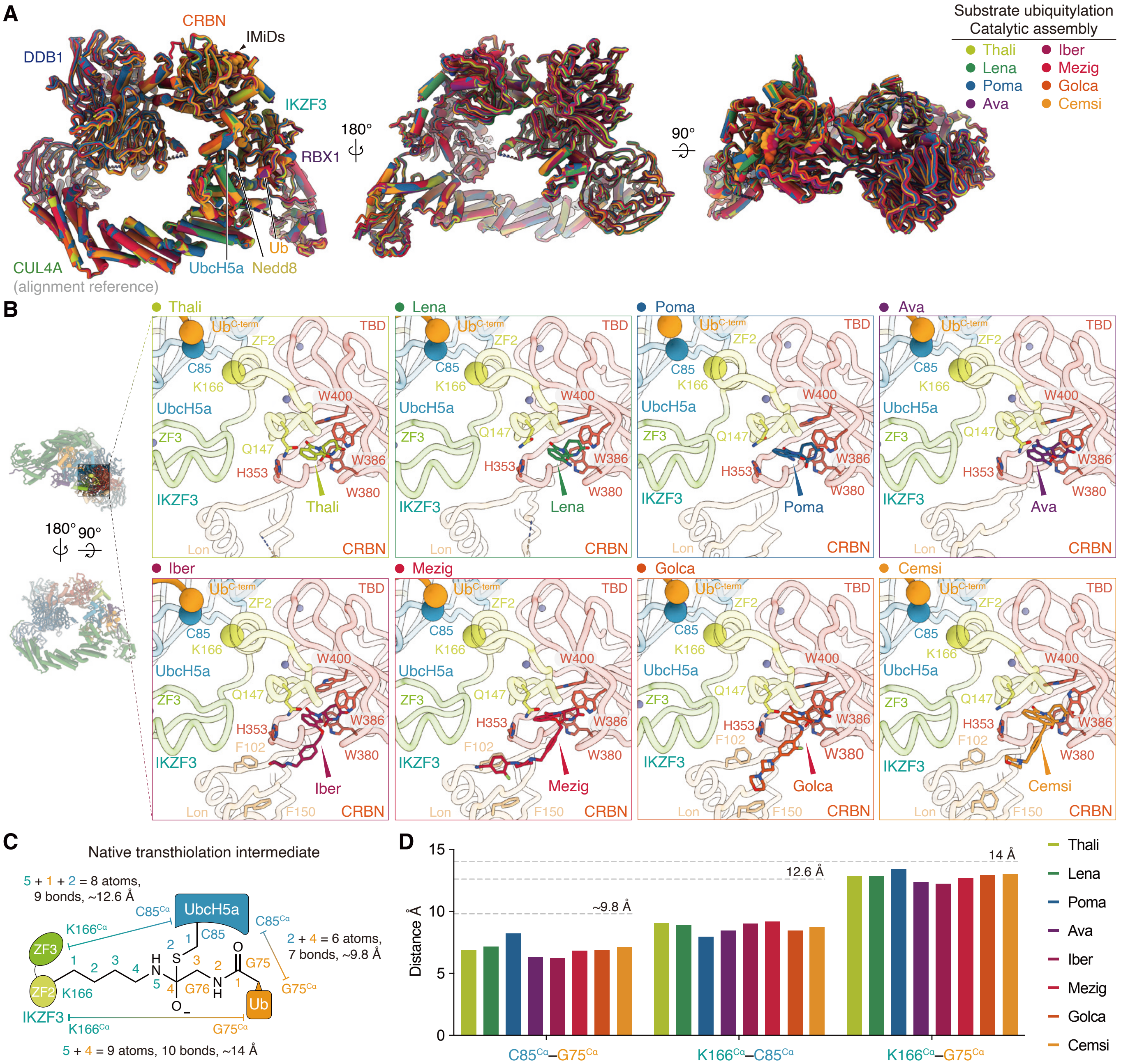
A
CRBN
IMiDs
DDB1
IKZF3
RBX1
180°
90°
CUL4A
(alignment reference)
UbcH5a
Nedd8
Ub
Substrate ubiquitylation
Catalytic assembly
Thali
Lena
Poma
Ava
Iber
Mezig
Golca
Cemsi
B
Thali
Lena
Poma
Ava
Iber
Mezig
Golca
Cemsi
Ub C-term
TBD
ZF2
C85
K166
UbcH5a
W400
ZF3
Q147
H353
W386
W380
IKZF3
F102
F150
Lon
CRBN
C
Native transthiolation intermediate
5 + 1 + 2 = 8 atoms,
9 bonds, ~12.6 Å
2 + 4 = 6 atoms,
7 bonds, ~9.8 Å
5 + 4 = 9 atoms, 10 bonds, ~14 Å
UbcH5a
C85Cα
K166Cα
G75Cα
ZF3
ZF2
IKZF3
K166
C85
G76
G75
Ub
D
Distance Å
~9.8 Å
12.6 Å
14 Å
C85Cα–G75Cα
K166Cα–C85Cα
K166Cα–G75Cα

**Fig. S6. The tight organization of the active ubiquitylation assembly.**

(**A**) Three orientations of superposition of eight IMiD-organized ubiquitylation assembly structures. Structures were aligned based on CUL4A. A guide to coloring the eight structures is shown on the right.

(**B**) Zoomed-in views of the IMiD binding region and substrate ubiquitylation region in eight IMiD-organized ubiquitylation assemblies. Side chains of key interacting residues on CRBN and IKZF3 are shown as sticks. The Cα atoms of ubiquitylation site IKZF3 K166, UbcH5a catalytic cysteine C85, and Ub G75 are shown as spheres.

(**C**) Chemical structure and geometry of the native transthiolation intermediate in IMiD-organized ubiquitylation assembly. The spatial distance between each crosslinking site was estimated based on the number of covalent bonds involved, multiplied by the theoretical bond length of a carbon–carbon single bond (~1.4 Å per bond). This approximation provides an upper-bound estimate of the spatial distance under the assumption of fully extended bond geometries.

(**D**) Bar graph showing the spatial distance between each crosslinking site measured across the eight IMiD-organized ubiquitylation assemblies. The upper-bound estimates of the spatial distance are indicated by gray dashed lines.

### Figure S7

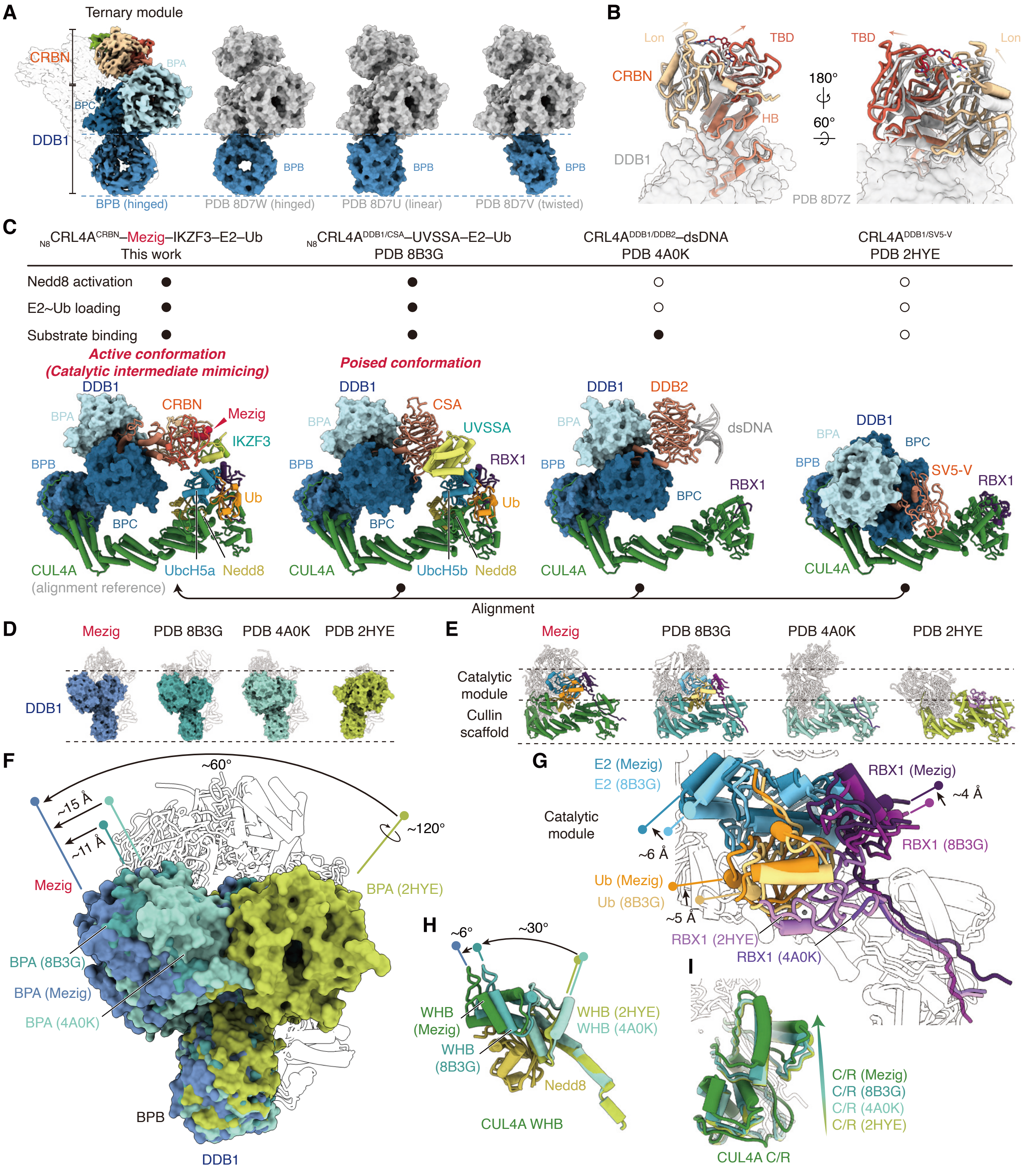
A
Ternary module
CRBN
BPA
BPC
DDB1
BPB
BPB (hinged)
PDB 8D7W (hinged)
PDB 8D7U (linear)
PDB 8D7V (twisted)
B
Lon
TBD
CRBN
HB
DDB1
180°
60°
PDB 8D7Z
C
N8CRL4ACRBN–Mezig–IKZF3–E2–Ub
This work
N8CRL4ADDB1/CSA–UVSSA–E2–Ub
PDB 8B3G
CRL4ADDB1/DDB2–dsDNA
PDB 4A0K
CRL4ADDB1/SV5-V
PDB 2HYE
Nedd8 activation
E2~Ub loading
Substrate binding
Active conformation
(Catalytic intermediate mimicing)
Poised conformation
DDB1
CRBN
Mezig
IKZF3
CSA
UVSSA
RBX1
DDB2
dsDNA
SV5-V
Ub
BPC
CUL4A
(alignment reference)
UbcH5a
UbcH5b
Nedd8
Alignment
D
Mezig
PDB 8B3G
PDB 4A0K
PDB 2HYE
DDB1
E
Catalytic module
Cullin scaffold
F
~60°
~15 Å
~11 Å
~120°
BPA (2HYE)
BPA (8B3G)
BPA (Mezig)
BPA (4A0K)
BPB
DDB1
G
E2 (Mezig)
E2 (8B3G)
RBX1 (Mezig)
~4 Å
RBX1 (8B3G)
~6 Å
Ub (Mezig)
Ub (8B3G)
~5 Å
RBX1 (2HYE)
RBX1 (4A0K)
H
~6°
~30°
WHB (Mezig)
WHB (8B3G)
WHB (2HYE)
WHB (4A0K)
Nedd8
CUL4A WHB
I
C/R (Mezig)
C/R (8B3G)
C/R (4A0K)
C/R (2HYE)
CUL4A C/R

**Fig. S7. Striking conformational flexibility of the $_{N8}$CRL4$^{CRBN}$ E3 ligase in catalyzing IMiD-induced neosubstrate ubiquitylation and different inactive states.**

(**A**) Cryo-EM map of CRBN/DDB1 in the ternary module within the mezigdomide-organized ubiquitylation assembly and surface presentations of reported mezigdomide-liganded CRBN/DDB1 in three discrete conformations (hinged, PDB 8D7W; linear, PDB 8D7U; twisted, PDB 8D7V) are shown from the same orientation. The CUL4A-interacting DDB1 BPB domain (colored blue) in the ubiquitylation assembly adopts the hinged conformation.

(**B**) Structural alignment of CRBN/DDB1 within the mezigdomide-organized ubiquitylation assembly and reported mezigdomide-liganded CRBN/DDB1 ternary complex (PDB 8D7Z). The substrates in these structures were hidden for clarity. Conformational changes are indicated by arrows.

(**C**) Description and cartoon models of four reported structures representing different states of the CRL4A E3 ligase complex. States were defined according to Nedd8 activation status, E2~Ub loading, and substrate binding. Structures were aligned to the Mezig structure based on CUL4A for further analysis in **D**–**I**.

(**D**) Surface presentations of DDB1 in the four structures are shown from the same orientation. Other components are contoured by black lines.

(**E**) Cartoon presentations of the catalytic module and/or cullin scaffold in the four structures are shown from the same orientation. Other components are contoured by black lines.

(**F**) Superposition of the four structures, highlighting the stepwise conformational changes of the DDB1 BPA domain.

(**G**) Superposition of the four structures, highlighting the stepwise conformational changes of the catalytic module containing RBX1 and E2~Ub.

(**H**) Superposition of the four structures, highlighting the stepwise conformational changes of the CUL4A WHB domain.

(**I**) Superposition of the four structures, highlighting the stepwise conformational changes of the CUL4A C/R domain.

### Figure S8

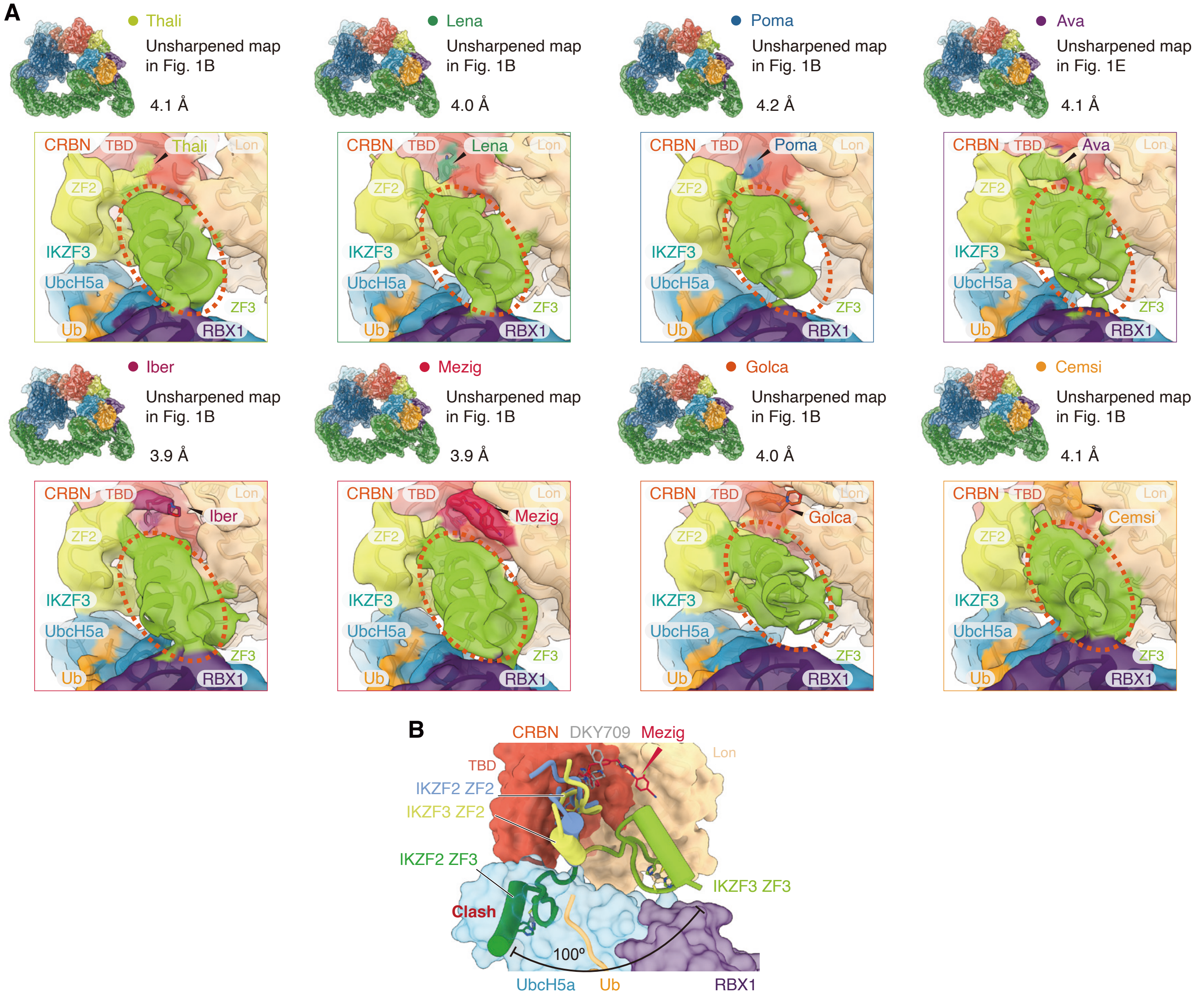
A
Thali
Unsharpened map in Fig. 1B
4.1 Å
Lena
Unsharpened map in Fig. 1B
4.0 Å
Poma
Unsharpened map in Fig. 1B
4.2 Å
Ava
Unsharpened map in Fig. 1E
4.1 Å
CRBN
TBD
Lon
ZF2
IKZF3
UbcH5a
ZF3
Ub
RBX1
Iber
Unsharpened map in Fig. 1B
3.9 Å
Mezig
Unsharpened map in Fig. 1B
3.9 Å
Golca
Unsharpened map in Fig. 1B
4.0 Å
Cemsi
Unsharpened map in Fig. 1B
4.1 Å
B
CRBN
DKY709
Mezig
TBD
Lon
IKZF2 ZF2
IKZF3 ZF2
IKZF2 ZF3
IKZF3 ZF3
Clash
100°
UbcH5a
Ub
RBX1

**Fig. S8. Unexpected cryo-EM density of IKZF3 ZF3 bridges the CRBN Lon domain and RBX1/UbcH5a.**

(**A**) Zoomed-in views of unsharpened cryo-EM maps of eight IMiD-organized ubiquitylation assemblies. Red dashed circles highlight densities of the IKZF3 ZF3 motif.

(**B**) Superposition of the mezigdomide-organized ubiquitylation assembly structure and the IKZF2–DKY709–CRBN/DDB1 ternary complex (PDB 8DEY) structure. Spatial differences in the positioning of the ZF3 motifs and potential steric clashes are indicated.

### Figure S9

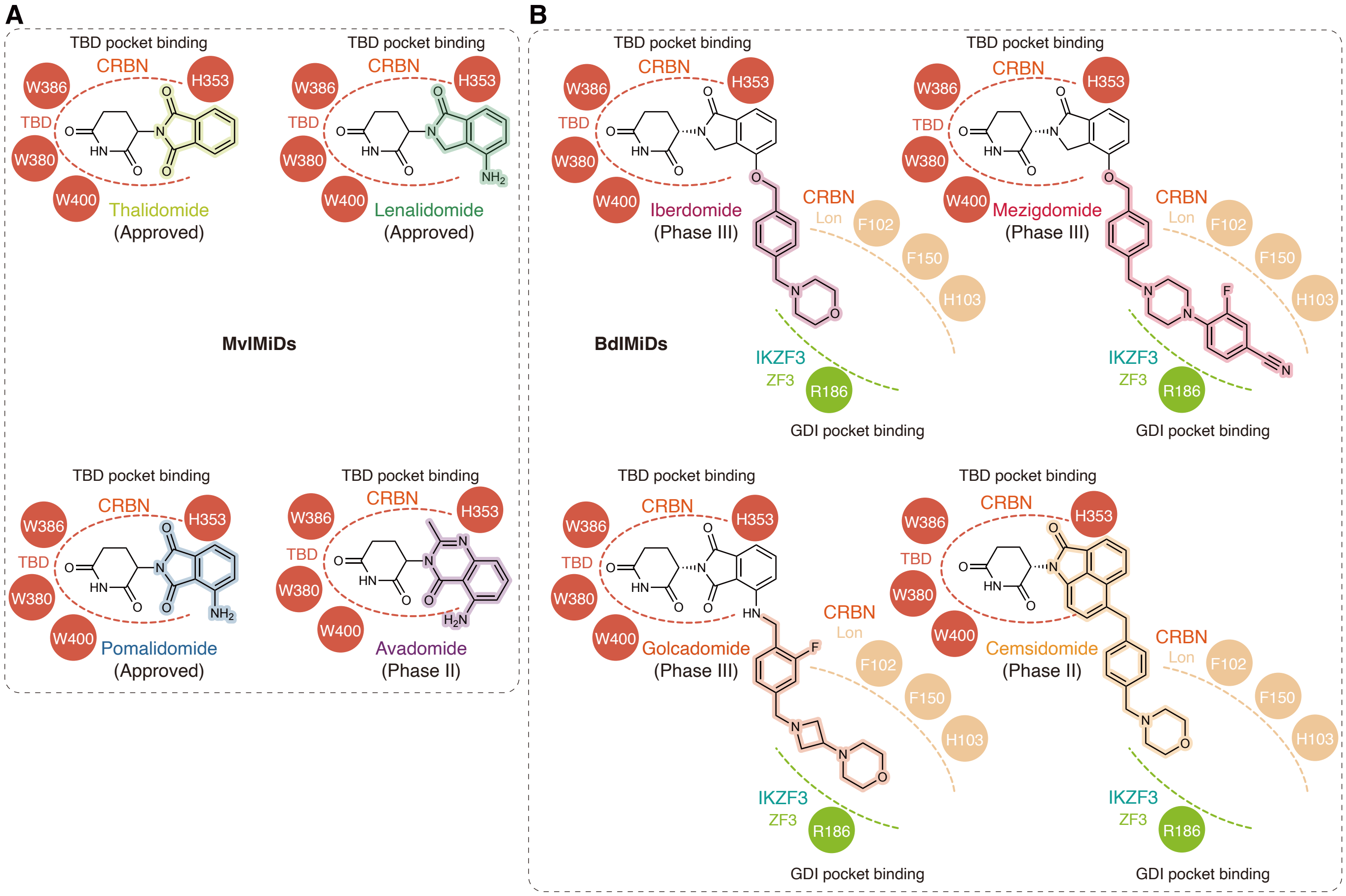
A
B
TBD pocket binding
CRBN
TBD
W386
H353
W380
W400
Thalidomide
(Approved)
Lenalidomide
(Approved)
MvIMiDs
Pomalidomide
(Approved)
Avadomide
(Phase II)
BdIMiDs
Iberdomide
(Phase III)
Mezigdomide
(Phase III)
Golcadomide
(Phase III)
Cemsidomide
(Phase II)
Lon
F102
F150
H103
IKZF3
ZF3
R186
GDI pocket binding

**Fig. S9. Comparison of IMiDs' chemical structures and interactions with CRBN/IKZF3.**

(**A**, **B**) Schematic IMiD–TBD pocket interactions and IMiD–GDI pocket interactions of MvIMiDs (**A**) or BdIMiDs (**B**).

### Figure S10

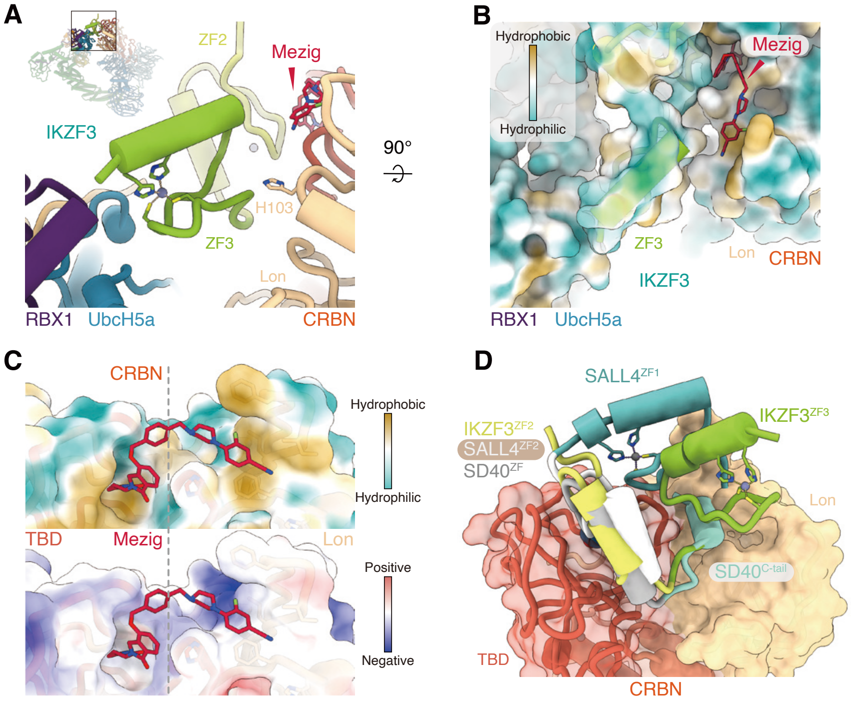
A
ZF2
Mezig
IKZF3
90°
H103
ZF3
Lon
RBX1
UbcH5a
CRBN
B
Hydrophobic
Hydrophilic
Mezig
ZF3
Lon
CRBN
IKZF3
RBX1
UbcH5a
C
CRBN
Hydrophobic
Hydrophilic
TBD
Mezig
Lon
Positive
Negative
D
SALL4ZF1
IKZF3ZF2
SALL4ZF2
SD40ZF
IKZF3ZF3
Lon
SD40C-tail
TBD
CRBN

**Fig. S10. CRBN Lon domain provides an interface for the synergistic binding of IMiDs and zinc finger proteins.**

(**A**) Detailed view shows IKZF3 ZF3 bridging the CRBN Lon domain and RBX1/UbcH5a.

(**B**) The hydrophobic/hydrophilic surface properties of IKZF3, CRBN, RBX1, and UbcH5a.

(**C**) The hydrophobic/hydrophilic properties (top) and electrostatic properties (bottom) of the CRBN TBD and Lon surface binding with mezigdomide.

(**D**) Structural alignments of the IKZF3–mezigdomide–CRBN/DDB1 ternary module (this work), the SALL4–iberdomide–CRBN/DDB1 ternary complex (PDB 8U16), and SD40–PT-179–CRBN/DDB1 ternary complex (PDB 8TNP), showing various zinc finger proteins binding to the CRBN Lon surface.

### Figure S11

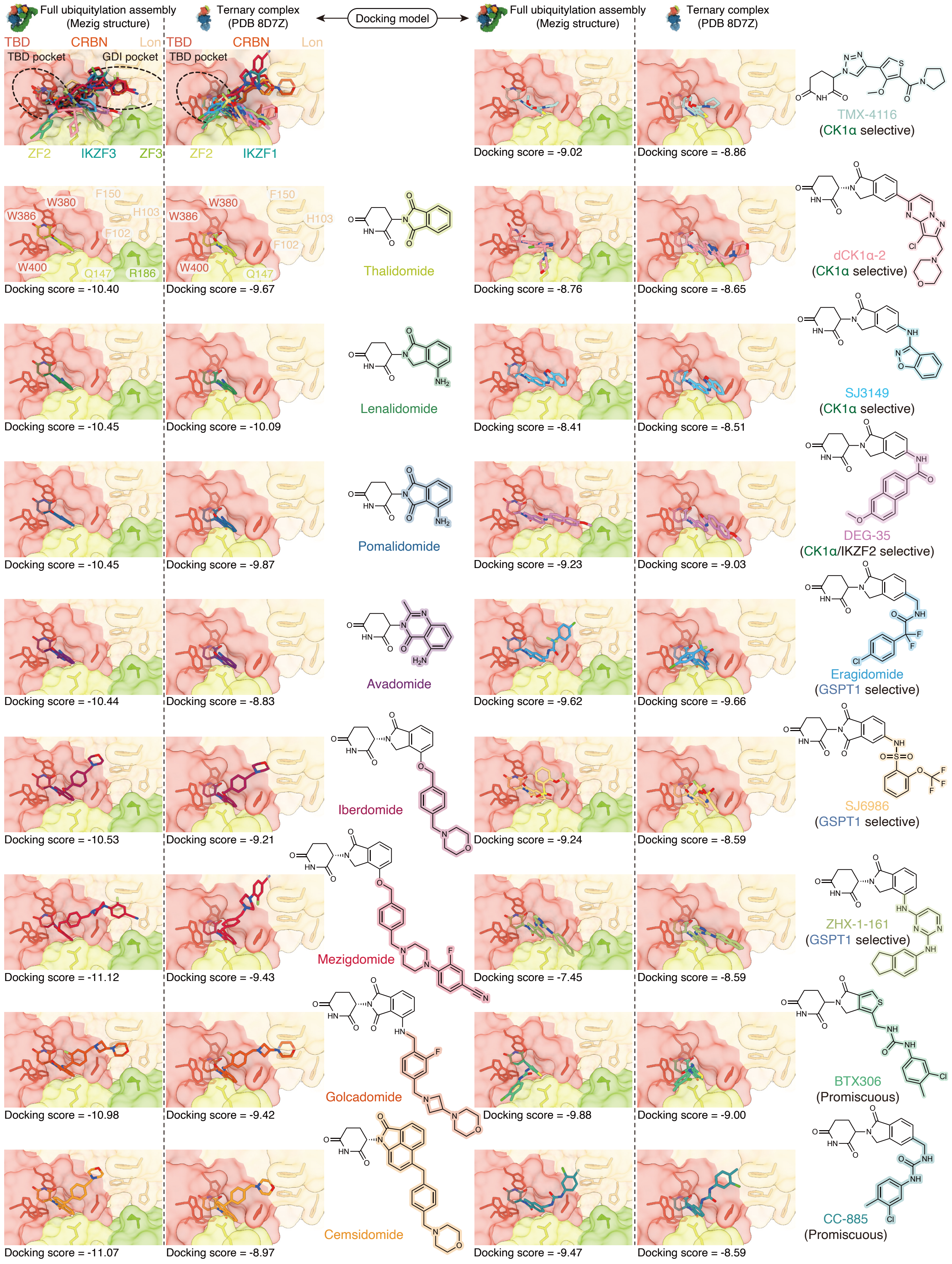

Full ubiquitylation assembly (Mezig structure)
Ternary complex (PDB 8D7Z)
Docking model
TBD
CRBN
Lon
TBD pocket
GDI pocket
ZF2
IKZF3
ZF3
IKZF1
W380
W386
F150
H103
F102
W400
Q147
R186
Thalidomide
Lenalidomide
Pomalidomide
Avadomide
Iberdomide
Mezigdomide
Golcadomide
Cemsidomide
TMX-4116
(CK1α selective)
dCK1α-2
(CK1α selective)
SJ3149
(CK1α selective)
DEG-35
(CK1α/IKZF2 selective)
Eragidomide
(GSPT1 selective)
SJ6986
(GSPT1 selective)
ZHX-1-161
(GSPT1 selective)
BTX306
(Promiscuous)
CC-885
(Promiscuous)
Docking score = -10.40
Docking score = -9.67
Docking score = -10.45
Docking score = -10.09
Docking score = -10.45
Docking score = -9.87
Docking score = -10.44
Docking score = -8.83
Docking score = -10.53
Docking score = -9.21
Docking score = -11.12
Docking score = -9.43
Docking score = -10.98
Docking score = -9.42
Docking score = -11.07
Docking score = -8.97
Docking score = -9.02
Docking score = -8.86
Docking score = -8.76
Docking score = -8.65
Docking score = -8.41
Docking score = -8.51
Docking score = -9.23
Docking score = -9.03
Docking score = -9.62
Docking score = -9.66
Docking score = -9.24
Docking score = -8.59
Docking score = -7.45
Docking score = -8.59
Docking score = -9.88
Docking score = -9.00
Docking score = -9.47
Docking score = -8.59

**Fig. S11. Docking poses of IMiDs with distinct substrate selectivities within the full ubiquitylation complex or ternary complex.**

Docking poses and docking scores of each IMiD molecule in the CRBN TBD pocket and/or GDI pocket. CRBN and IKZF1/3 are shown as surface representations. Side chains of key CRBN and IKZF1/3 interacting residues, including CRBN H103, F102, F150, H353, W380, W386, W400, IKZF3 Q147 (IKZF1 Q146), and IKZF3 R186 are indicated. Chemical structures of each IMiD molecule are shown beside the docking model.

### Figure S12

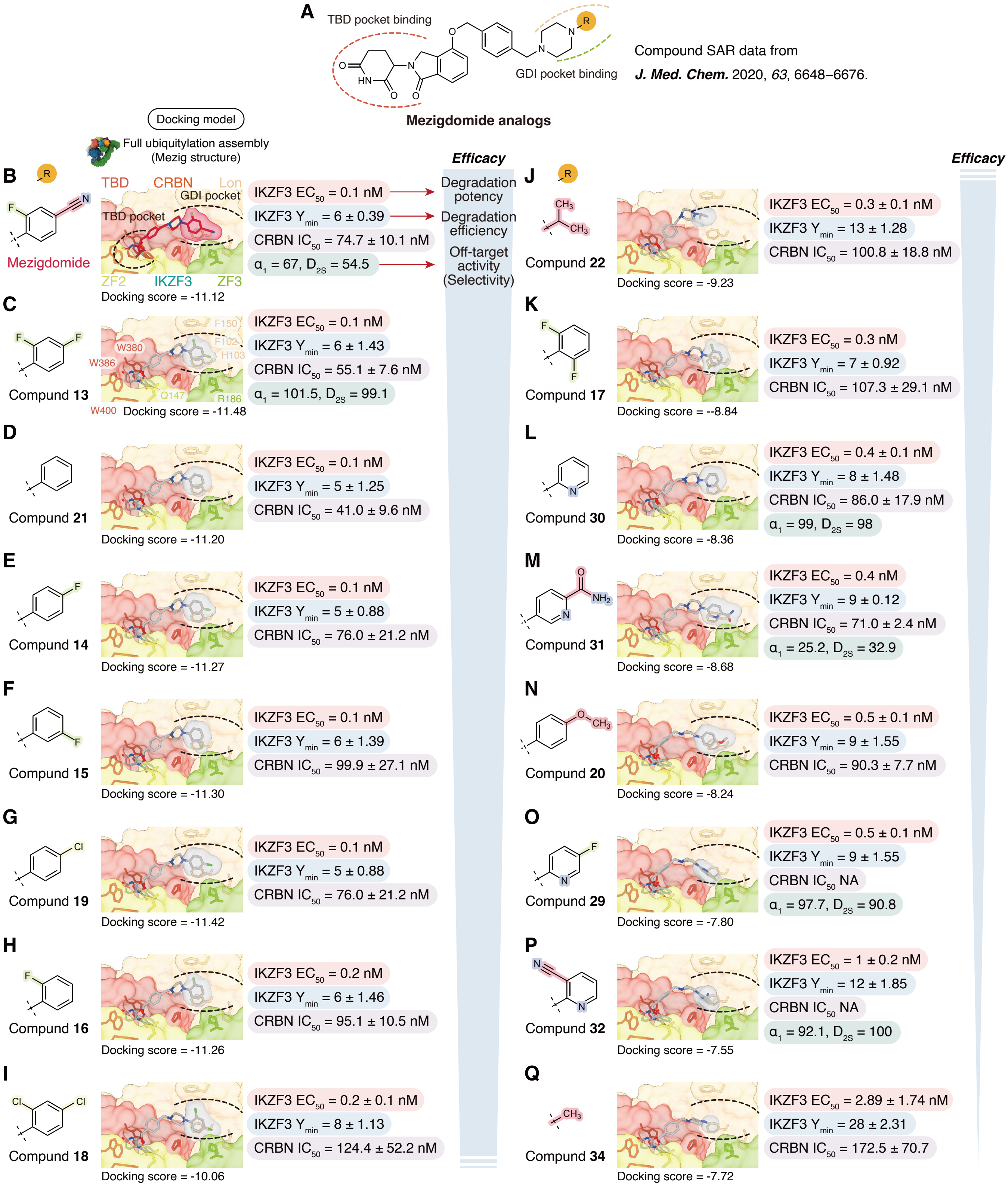
A
TBD pocket binding
GDI pocket binding
Mezigdomide analogs
Compound SAR data from
J. Med. Chem. 2020, 63, 6648–6676.
Docking model
Full ubiquitylation assembly (Mezig structure)
Efficacy
B
Mezigdomide
TBD
CRBN
Lon
GDI pocket
TBD pocket
ZF2
IKZF3
ZF3
Docking score = -11.12
IKZF3 EC50 = 0.1 nM
IKZF3 Ymin = 6 ± 0.39
CRBN IC50 = 74.7 ± 10.1 nM
α1 = 67, D2S = 54.5
Degradation potency
Degradation efficiency
Off-target activity (Selectivity)
C
Compund 13
F150
F102
H103
W380
W386
Q147
R186
W400
Docking score = -11.48
IKZF3 EC50 = 0.1 nM
IKZF3 Ymin = 6 ± 1.43
CRBN IC50 = 55.1 ± 7.6 nM
α1 = 101.5, D2S = 99.1
D
Compund 21
Docking score = -11.20
IKZF3 EC50 = 0.1 nM
IKZF3 Ymin = 5 ± 1.25
CRBN IC50 = 41.0 ± 9.6 nM
E
Compund 14
Docking score = -11.27
IKZF3 EC50 = 0.1 nM
IKZF3 Ymin = 5 ± 0.88
CRBN IC50 = 76.0 ± 21.2 nM
F
Compund 15
Docking score = -11.30
IKZF3 EC50 = 0.1 nM
IKZF3 Ymin = 6 ± 1.39
CRBN IC50 = 99.9 ± 27.1 nM
G
Compund 19
Docking score = -11.42
IKZF3 EC50 = 0.1 nM
IKZF3 Ymin = 5 ± 0.88
CRBN IC50 = 76.0 ± 21.2 nM
H
Compund 16
Docking score = -11.26
IKZF3 EC50 = 0.2 nM
IKZF3 Ymin = 6 ± 1.46
CRBN IC50 = 95.1 ± 10.5 nM
I
Compund 18
Docking score = -10.06
IKZF3 EC50 = 0.2 ± 0.1 nM
IKZF3 Ymin = 8 ± 1.13
CRBN IC50 = 124.4 ± 52.2 nM
Efficacy
J
Compund 22
Docking score = -9.23
IKZF3 EC50 = 0.3 ± 0.1 nM
IKZF3 Ymin = 13 ± 1.28
CRBN IC50 = 100.8 ± 18.8 nM
K
Compund 17
Docking score = --8.84
IKZF3 EC50 = 0.3 nM
IKZF3 Ymin = 7 ± 0.92
CRBN IC50 = 107.3 ± 29.1 nM
L
Compund 30
Docking score = -8.36
IKZF3 EC50 = 0.4 ± 0.1 nM
IKZF3 Ymin = 8 ± 1.48
CRBN IC50 = 86.0 ± 17.9 nM
α1 = 99, D2S = 98
M
Compund 31
Docking score = -8.68
IKZF3 EC50 = 0.4 nM
IKZF3 Ymin = 9 ± 0.12
CRBN IC50 = 71.0 ± 2.4 nM
α1 = 25.2, D2S = 32.9
N
Compund 20
Docking score = -8.24
IKZF3 EC50 = 0.5 ± 0.1 nM
IKZF3 Ymin = 9 ± 1.55
CRBN IC50 = 90.3 ± 7.7 nM
O
Compund 29
Docking score = -7.80
IKZF3 EC50 = 0.5 ± 0.1 nM
IKZF3 Ymin = 9 ± 1.55
CRBN IC50 NA
α1 = 97.7, D2S = 90.8
P
Compund 32
Docking score = -7.55
IKZF3 EC50 = 1 ± 0.2 nM
IKZF3 Ymin = 12 ± 1.85
CRBN IC50 NA
α1 = 92.1, D2S = 100
Q
Compund 34
Docking score = -7.72
IKZF3 EC50 = 2.89 ± 1.74 nM
IKZF3 Ymin = 28 ± 2.31
CRBN IC50 = 172.5 ± 70.7

**Fig. S12. Understanding the structure-activity relationship (SAR) of mezigdomide analogs through GDI pocket interacting models.**

(**A**) Chemical structure of mezigdomide analogs. The terminal functional group conjugated to the piperizine moiety is indicated by the "R" group.

(**B**–**Q**) The chemical structure of the R group in each mezigdomide analog (left), the docking model and docking scores of the analog within the full ubiquitylation complex (middle), and reported SAR data (right). For SAR data, IKZF3 EC50 indicates the degradation potency. IKZF3 $Y_{min}$, the protein remaining after drug treatment, indicates the degradation efficiency. Value of $\alpha_1$ and $D_{2S}$ mean percent inhibition of $\alpha_1$ adrenergic receptor and dopamine $D_{2S}$ receptor, which indicates off-target activity (or selectivity). The reported compounds are ranked from high to low efficacy, as indicated by the light blue arrow. The most striking difference arises from the comparison between compound **34** (**Q**) and other analogs: replacing the terminal aromatic ring with a smaller methyl group significantly reduces compatibility with the GDI pocket, accompanied by a marked impairment in IKZF3 EC50, $Y_{min}$, and CRBN IC50 values. The substitution of the 4-nitrile group (**B**) with a fluorine atom (**C**) compromises the fit within the GDI pocket, which is associated with a notable decline in the selectivity of the drug.

**Table S1. Cryo-EM data collection, refinement, and validation statistics for MvIMiD structures.**

| | $_{N8}$CRL4$^{CRBN}$–IMiD–IKZF3(ZF2–ZF3)–UbcH5a–Ub complex | | | |
|---|---|---|---|---|
| **MvIMiDs** | Thalidomide | Lenalidomide | Pomalidomide | Avadomide |
| **PDB entry** | 9UUQ | 9V09 | 9V0A | 9V0B |
| **EMDB entry** | EMD-64515 | EMD-64658 | EMD-64659 | EMD-64660 |
| **Data collection & processing** | | | | |
| **Microscope and detector** | Krios G4/Falcon4 | Krios G4/Falcon4 | Krios/K3 | Krios G4/Falcon4 |
| **Magnification** | 75,000 | 75,000 | 81,000 | 75,000 |
| **Voltage (keV)** | 300 | 300 | 300 | 300 |
| **Electron exposure ($e^-$/Å$^2$)** | 50 | 50 | 50 | 50 |
| **Defocus range (µm)** | –1.2 to –2.0 | –1.2 to –2.0 | –1.0 to –1.8 | –1.2 to –2.0 |
| **Pixel size (Å)** | 1.059 | 1.059 | 1.074 | 1.059 |
| **Symmetry imposed** | C1 | C1 | C1 | C1 |
| **Micrographs (no.)** | 7,706 | 7,621 | 13,668 | 8,474 |
| **Final particles (no.)** | 62,925 | 89,694 | 180,420 | 60,674 |
| **Map global resolution (Å)** | 3.61 | 3.51 | 3.82 | 3.61 |
| **FSC threshold** | 0.143 | 0.143 | 0.143 | 0.143 |
| **$CC_{mask}$/$CC_{volume}$** | 0.78/0.77 | 0.77/0.76 | 0.81/0.81 | 0.76/0.74 |
| **B-factor (Å$^2$)** | –70 | –70 | –100 | –80 |
| **Refinement & validation** | | | | |
| **Initial model** | PDB 8B3G, 8D7Z; AF-Q9UKT9-F1-v4 | PDB 8B3G, 8D7Z; AF-Q9UKT9-F1-v4 | PDB 8B3G, 8D7Z; AF-Q9UKT9-F1-v4 | PDB 8B3G, 8D7Z; AF-Q9UKT9-F1-v4 |
| **Model composition** | | | | |
| **Non-hydrogen atoms** | 20,534 | 20,551 | 20,824 | 20,732 |
| **Protein residues** | 2,581 | 2,586 | 2,616 | 2,614 |
| **Ligands** | 6 Zn, 1 Thali | 6 Zn, 1 Lena | 6 Zn, 1 Poma | 6 Zn, 1 Ava |
| **R.M.S. deviations** | | | | |
| **Bond lengths (Å)** | 0.003 | 0.003 | 0.004 | 0.003 |
| **Bond angles (°)** | 0.682 | 0.669 | 0.699 | 0.660 |
| **B-factor (Å$^2$)** | | | | |
| **Protein (min/max/mean)** | 30.00/246.66/102.14 | 21.23/323.37/89.54 | 30.00/324.79/137.67 | 17.58/240.23/84.91 |
| **Ligand (min/max/mean)** | 81.62/243.30/95.94 | 90.92/328.46/108.60 | 115.58/297.18/143.28 | 67.33/237.36/82.61 |
| **MolProbity score** | 1.78 | 1.80 | 1.92 | 1.78 |
| **Clashscore** | 7.10 | 7.54 | 10.53 | 7.02 |
| **Rotamer outliers (%)** | 0.13 | 0.04 | 0.04 | 0 |
| **CaBLAM outliers (%)** | 2.62 | 2.69 | 3.01 | 3.32 |
| **Ramachandran plot** | | | | |
| **Favored (%)** | 94.31 | 94.37 | 94.63 | 94.28 |
| **Allowed (%)** | 5.69 | 5.63 | 5.37 | 5.72 |
| **Outliers (%)** | 0 | 0 | 0 | 0 |

**Table S2. Cryo-EM data collection, refinement, and validation statistics for BdIMiD structures.**

| | $_{N8}$CRL4$^{CRBN}$–IMiD–IKZF3(ZF2–ZF3)–UbcH5a–Ub complex | | | |
|---|---|---|---|---|
| **BdIMiDs** | Iberdomide | Mezigdomide | Golcadomide | Cemsidomide |
| **PDB entry** | 9V0C | 9UUM | 9V0E | 9V0F |
| **EMDB entry** | EMD-64661 | EMD-64512 | EMD-64662 | EMD-64663 |
| **Data collection & processing** | | | | |
| **Microscope and detector** | Krios G4/Falcon4 | Krios G4/Falcon4 | Krios G4/Falcon4 | Krios G4/Falcon4 |
| **Magnification** | 75,000 | 96,000 | 75,000 | 75,000 |
| **Voltage (keV)** | 300 | 300 | 300 | 300 |
| **Electron exposure ($e^-$/Å$^2$)** | 50 | 50 | 50 | 50 |
| **Defocus range (µm)** | –1.2 to –2.0 | –1.2 to –2.0 | –1.2 to –2.0 | –1.2 to –2.0 |
| **Pixel size (Å)** | 1.059 | 0.83 | 1.059 | 1.059 |
| **Symmetry imposed** | C1 | C1 | C1 | C1 |
| **Micrographs (no.)** | 8,097 | 25,882 | 8,567 | 7,205 |
| **Final particles (no.)** | 42,369 | 172,162 | 37,205 | 48,755 |
| **Map global resolution (Å)** | 3.46 | 3.60 | 3.60 | 3.71 |
| **FSC threshold** | 0.143 | 0.143 | 0.143 | 0.143 |
| **$CC_{mask}$/$CC_{volume}$** | 0.79/0.78 | 0.81/0.78 | 0.70/0.69 | 0.77/0.76 |
| **B-factor (Å$^2$)** | –70 | –100 | –80 | –50 |
| **Refinement & validation** | | | | |
| **Initial model** | PDB 8B3G, 8D7Z; AF-Q9UKT9-F1-v4 | PDB 8B3G, 8D7Z; AF-Q9UKT9-F1-v4 | PDB 8B3G, 8D7Z; AF-Q9UKT9-F1-v4 | PDB 8B3G, 8D7Z; AF-Q9UKT9-F1-v4 |
| **Model composition** | | | | |
| **Non-hydrogen atoms** | 20,746 | 20,808 | 20,776 | 20,720 |
| **Protein residues** | 2,614 | 2,620 | 2,614 | 2,615 |
| **Ligands** | 6 Zn, 1 Iberdomide | 6 Zn, 1 Mezigdomide | 6 Zn, 1 Golcadomide | 6 Zn, 1 Cemsidomide |
| **R.M.S. deviations** | | | | |
| **Bond lengths (Å)** | 0.003 | 0.003 | 0.003 | 0.003 |
| **Bond angles (°)** | 0.669 | 0.626 | 0.569 | 0.695 |
| **B-factor (Å$^2$)** | | | | |
| **Protein (min/max/mean)** | 16.82/318.87/108.30 | 1.61/337.60/73.75 | 23.57/217.01/88.36 | 47.29/275.76/107.96 |
| **Ligand (min/max/mean)** | 84.78/234.61/116.96 | 63.10/171.33/70.48 | 34.39/133.74/89.09 | 108.61/311.36/120.29 |
| **MolProbity score** | 1.77 | 1.77 | 1.82 | 1.90 |
| **Clashscore** | 7.11 | 6.85 | 8.17 | 7.65 |
| **Rotamer outliers (%)** | 0 | 0.09 | 0 | 0 |
| **CaBLAM outliers (%)** | 2.58 | 2.77 | 3.01 | 3.24 |
| **Ramachandran plot** | | | | |
| **Favored (%)** | 94.43 | 94.25 | 94.51 | 92.35 |
| **Allowed (%)** | 5.57 | 5.75 | 5.45 | 7.65 |
| **Outliers (%)** | 0 | 0 | 0 | 0 |

**Table S3. Summary of reported binary or ternary complex structures in the PDB database that contain the eight IKZF1/3-targeting IMiDs examined in this study.**

| IMiDs | Number of binary complex | PDB Accession codes | Number of ternary complex | PDB Accession codes |
|---|---|---|---|---|
| **Thali** | 23 | 4CI1, 4TZC, 4V2Y, 4V32, 5AMH, 5AMI, 5AMJ, 5AMK, 5OH1, 5YIZ, 5YJ0, 6R0S, 6R0U, 6R12, 6R13, 6R18, 6R19, 6R1A, 6R1C, 6R1W 7PSO, 8BC7, 8OU3 | 1 | 7BQU |
| **Lena** | 5 | 4CI2, 4TZ4, 4V30, 9FJX, 8RQA | 1 | 5FQD |
| **Poma** | 5 | 4CI3, 4TZU, 4V2Z, 8D81, 8OIZ, | 7 | 6H0F, 6H0G, 6UML,8TNP, 8U15, 8U16, 8U17 |
| **Ava** | 1 | 7PSO | 0 | |
| **Iber** | 2 | 5V3O, 7PS9 | 2 | 8D80, 8U15 |
| **Mezig** | 4 | 8RQ8, 8D7U, 8D7V, 8D7W | 2 | 8RQC，8D7Z |
| **Golca** | 0 | -- | 0 | -- |
| **Cemsi** | 0 | -- | 0 | -- |

Data as of March 20, 2025.

**References**


1. Z. Deng *et al.*, Mechanistic insights into nucleosomal H2B monoubiquitylation mediated by yeast Bre1-Rad6 and its human homolog RNF20/RNF40-hRAD6A. *Mol. Cell* **83**, 3080-3094.e3014 (2023).
2. S. Diaz, L. Li, K. Wang, X. Liu, Expression and purification of functional recombinant CUL2•RBX1 from E. coli. *Sci. Rep.* **11**, 11224 (2021).
3. M. Pan *et al.*, Structural insights into Ubr1-mediated N-degron polyubiquitination. *Nature* **600**, 334-338 (2021).
4. J. Li *et al.*, Cullin-RING ligases employ geometrically optimized catalytic partners for substrate targeting. *Mol Cell* **84**, 1304-1320 e1316 (2024).
5. J. Liwocha *et al.*, Mechanism of millisecond Lys48-linked poly-ubiquitin chain formation by cullin-RING ligases. *Nat Struct Mol Biol* **31**, 378-389 (2024).
6. H. S. Ai *et al.*, Mechanism of nucleosomal H2A K13/15 monoubiquitination and adjacent dual monoubiquitination by RNF168. *Nat. Chem. Biol.*, (2024).
7. J. Lei, J. Frank, Automated acquisition of cryo-electron micrographs for single particle reconstruction on an FEI Tecnai electron microscope. *J. Struct. Biol.* **150**, 69-80 (2005).
8. J. Zivanov *et al.*, New tools for automated high-resolution cryo-EM structure determination in RELION-3. *eLife* **7**, e42166 (2018).
9. G. Kokic *et al.*, Structural basis for RNA polymerase II ubiquitylation and inactivation in transcription-coupled repair. *Nat. Struct. Mol. Biol.* **31**, (2024).
10. E. R. Watson *et al.*, Molecular glue CELMoD compounds are regulators of cereblon conformation. *Science* **378**, 549-553 (2022).
11. J. Jumper *et al.*, Highly accurate protein structure prediction with AlphaFold. *Nature* **596**, 583-589 (2021).
12. E. F. Pettersen *et al.*, UCSF chimera - A visualization system for exploratory research and analysis. *J. Comput. Chem.* **25**, 1605-1612 (2004).
13. P. Emsley, B. Lohkamp, W. G. Scott, K. Cowtan, Features and development of Coot. *Acta Crystallogr. D* **66**, 486-501 (2010).
14. P. D. Adams *et al.*, PHENIX: a comprehensive Python-based system for macromolecular structure solution. *Acta Crystallogr. D* **66**, 213-221 (2010).
15. E. S. Fischer *et al.*, Structure of the DDB1-CRBN E3 ubiquitin ligase in complex with thalidomide. *Nature* **512**, 49-53 (2014).
16. C. Heim, M. D. Hartmann, High-resolution structures of the bound effectors avadomide (CC-122) and iberdomide (CC-220) highlight advantages and limitations of the MsCI4 soaking system. *Acta Crystallographica Section D* **78**, 290-298 (2022).

#### Data S1-1

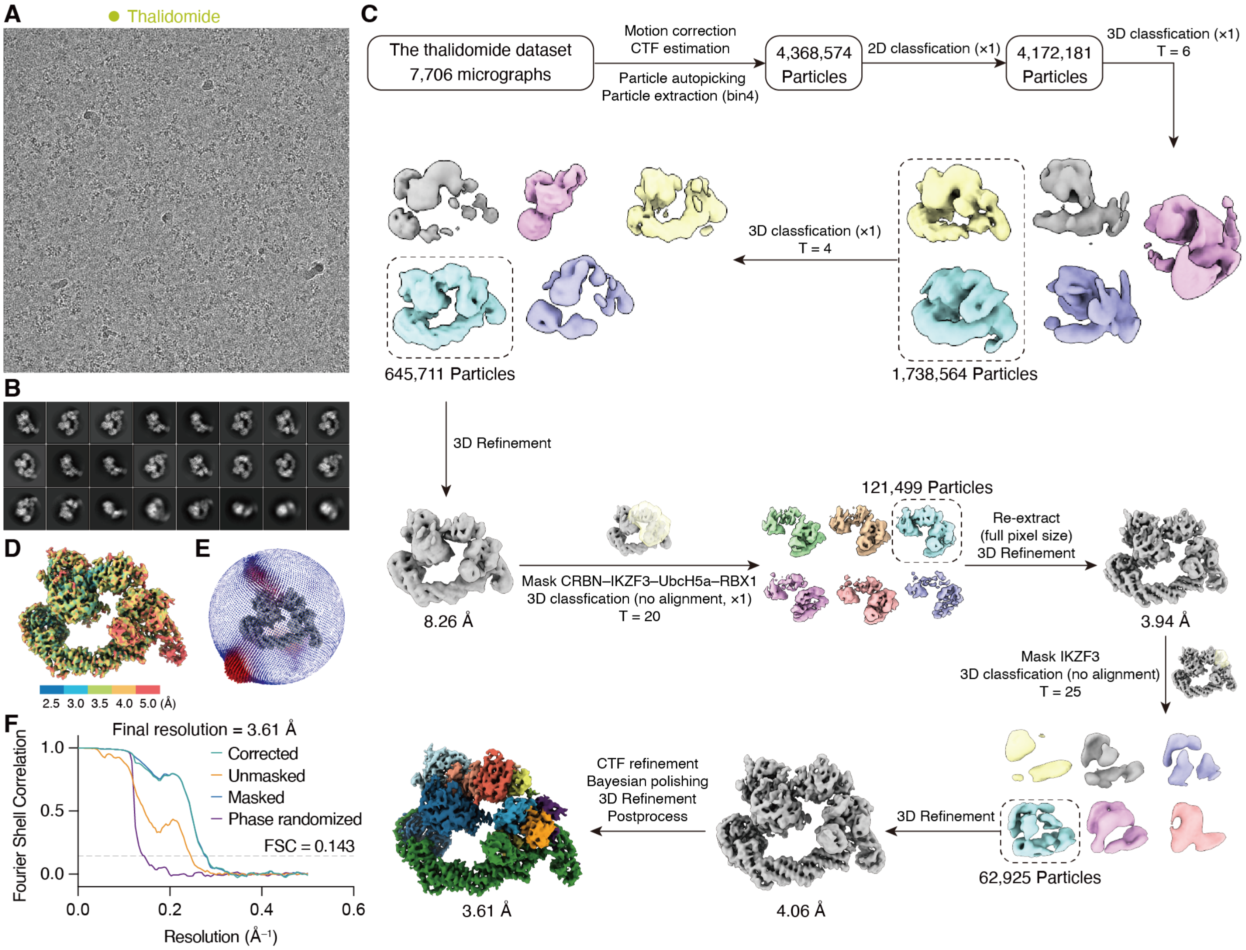


**Data S1-1. Cryo-EM data processing flow chart of the thalidomide-organized ubiquitylation assembly.** (**A**) A representative micrograph. (**B**) Representative 2D averages. (**C**) Data processing workflow. (**D**) Final cryo-EM map colored by the indicated local resolution. (**E**) Angular distributions of the final cryo-EM reconstruction. (**F**) Gold standard Fourier shell correlation (FSC) curves for the final refinement (3.61 Å resolution at FSC = 0.143).

#### Data S1-2

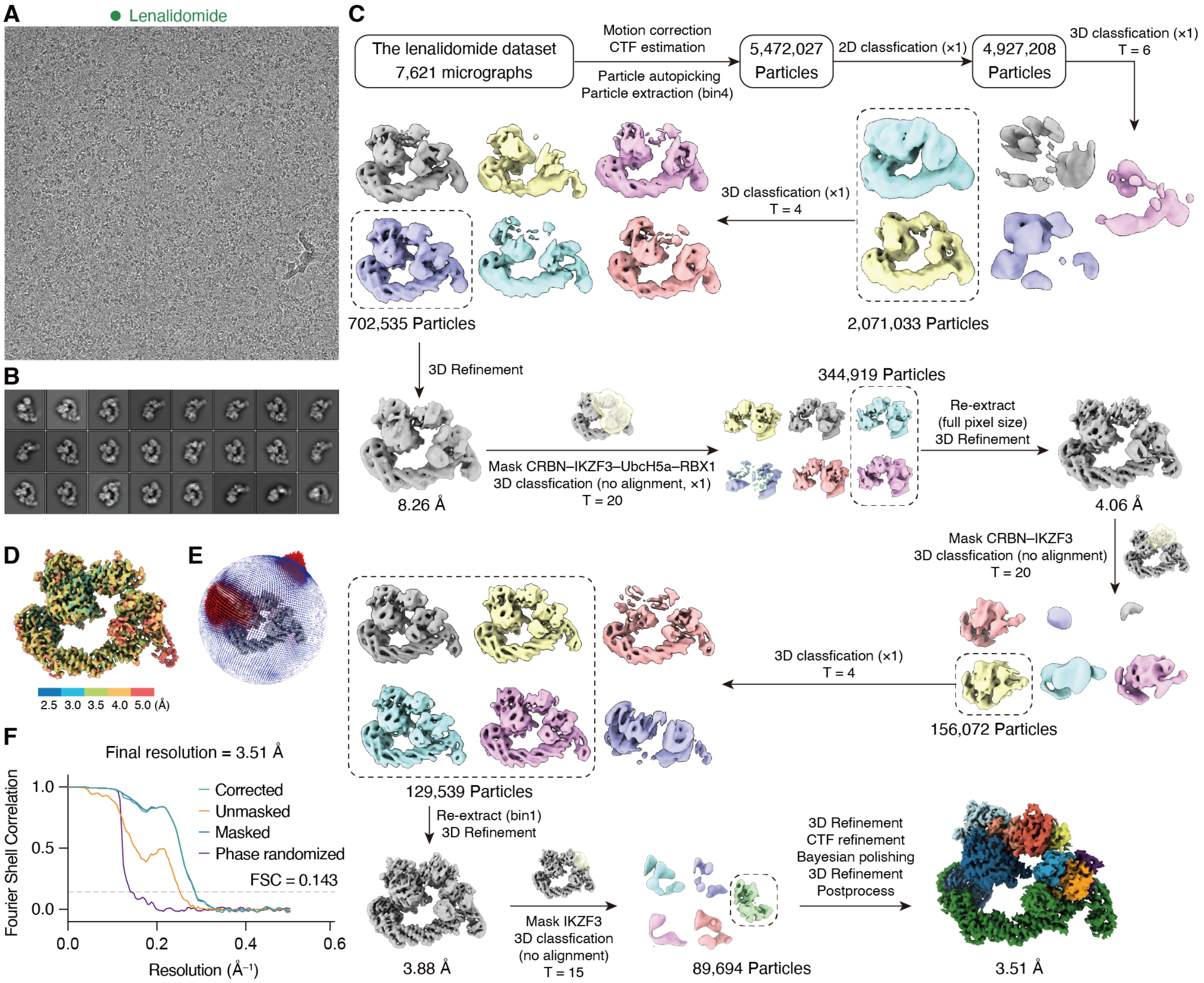


**Data S1-2. Cryo-EM data processing flow chart of the lenalidomide-organized ubiquitylation assembly.** (**A**) A representative micrograph. (**B**) Representative 2D averages. (**C**) Data processing workflow. (**D**) Final cryo-EM map colored by the indicated local resolution. (**E**) Angular distributions of the final cryo-EM reconstruction. (**F**) Gold standard Fourier shell correlation (FSC) curves for the final refinement (3.51 Å resolution at FSC = 0.143).

#### Data S1-3

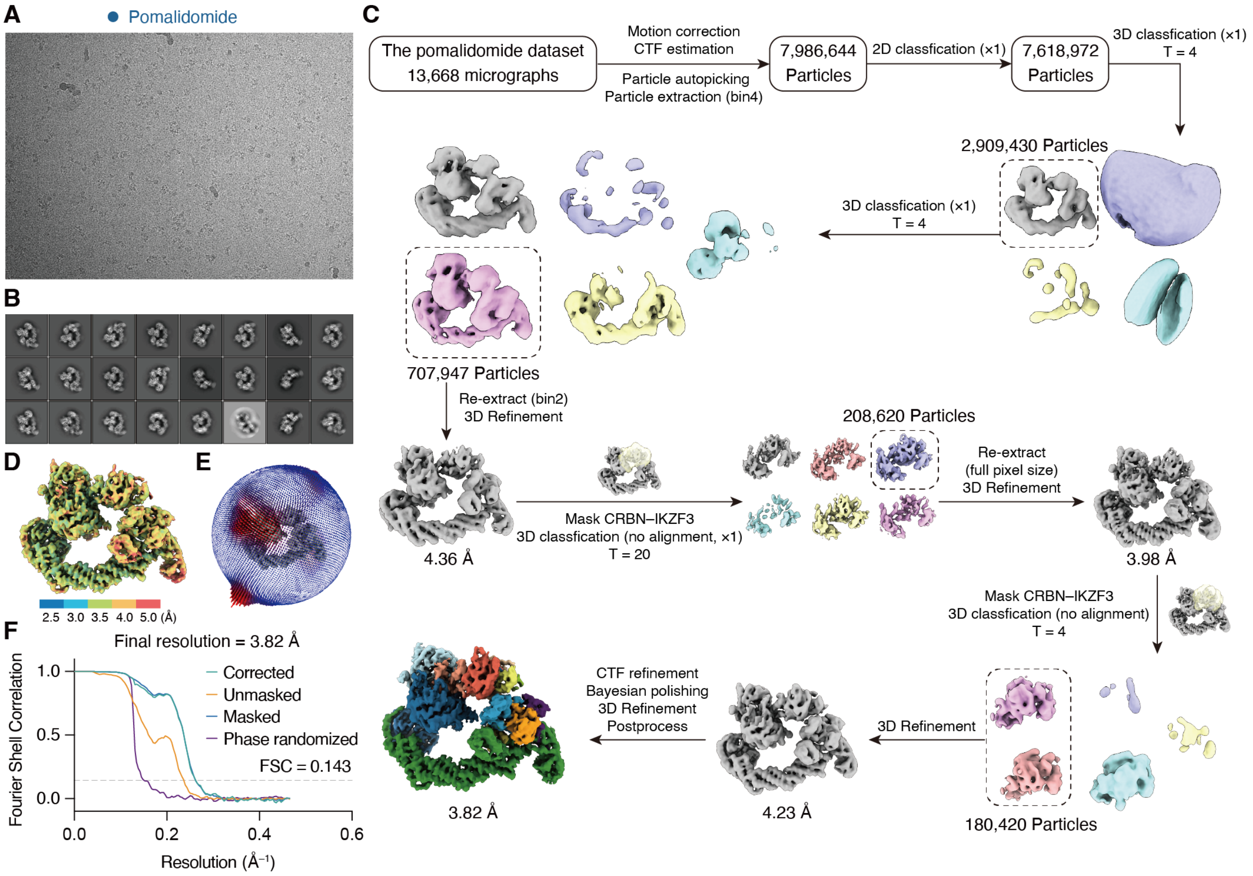


**Data S1-3. Cryo-EM data processing flow chart of the pomalidomide-organized ubiquitylation assembly.** (**A**) A representative micrograph. (**B**) Representative 2D averages. (**C**) Data processing workflow. (**D**) Final cryo-EM map colored by the indicated local resolution. (**E**) Angular distributions of the final cryo-EM reconstruction. (**F**) Gold standard Fourier shell correlation (FSC) curves for the final refinement (3.82 Å resolution at FSC = 0.143).

#### Data S1-4

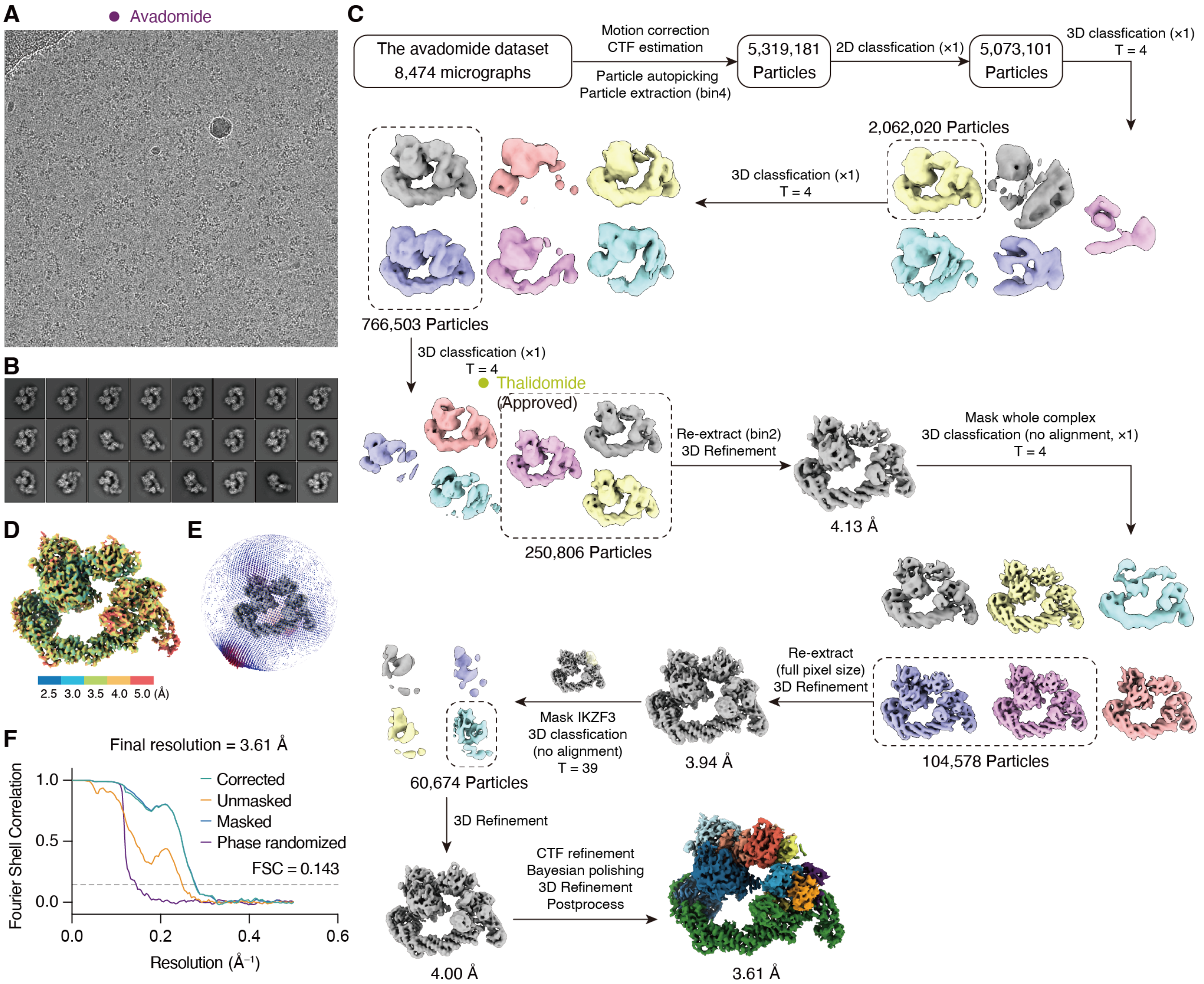


**Data S1-4. Cryo-EM data processing flow chart of the avadomide-organized ubiquitylation assembly.** (**A**) A representative micrograph. (**B**) Representative 2D averages. (**C**) Data processing workflow. (**D**) Final cryo-EM map colored by the indicated local resolution. (**E**) Angular distributions of the final cryo-EM reconstruction. (**F**) Gold standard Fourier shell correlation (FSC) curves for the final refinement (3.61 Å resolution at FSC = 0.143).

#### Data S1-5

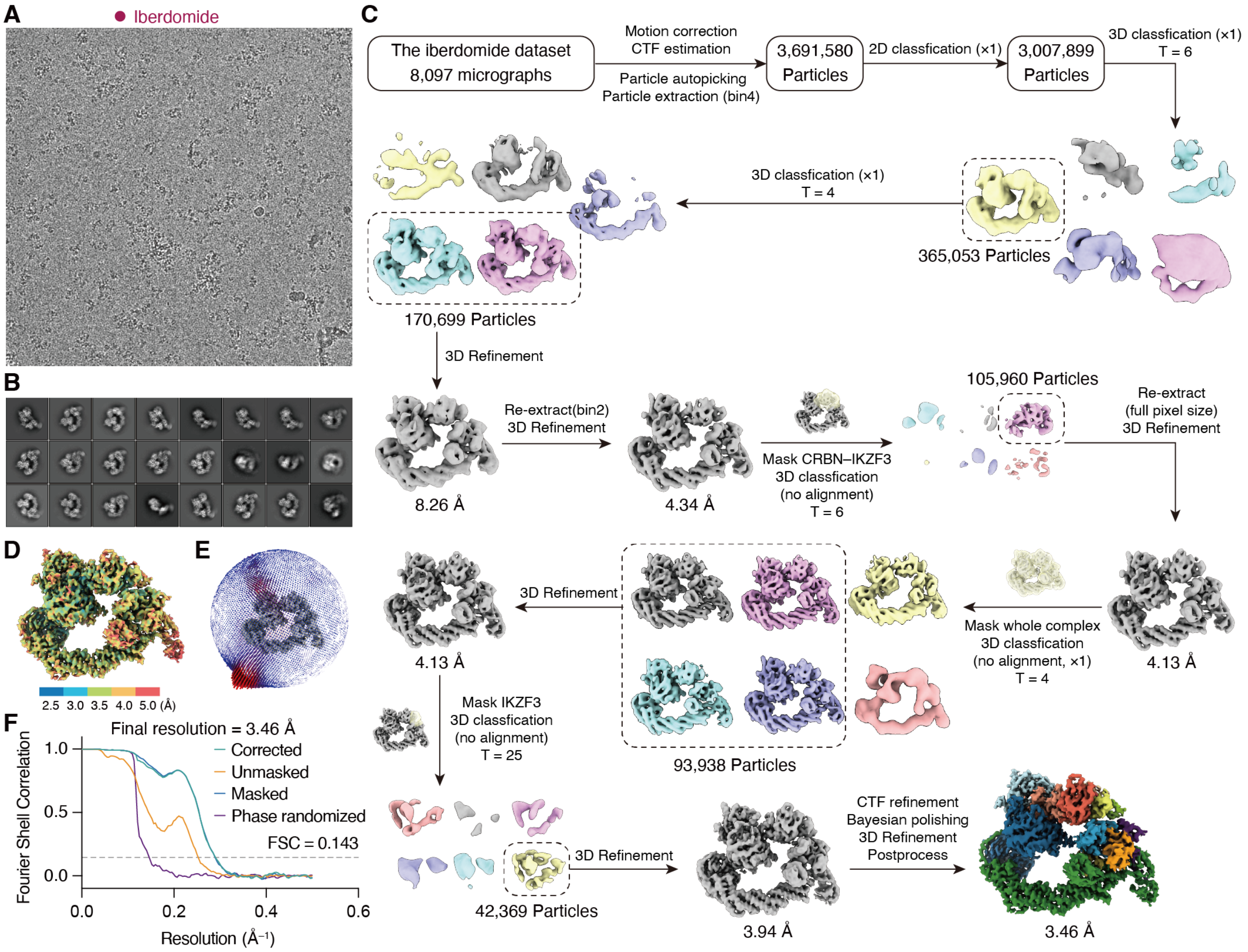


**Data S1-5. Cryo-EM data processing flow chart of the iberdomide-organized ubiquitylation assembly.** (**A**) A representative micrograph. (**B**) Representative 2D averages. (**C**) Data processing workflow. (**D**) Final cryo-EM map colored by the indicated local resolution. (**E**) Angular distributions of the final cryo-EM reconstruction. (**F**) Gold standard Fourier shell correlation (FSC) curves for the final refinement (3.46 Å resolution at FSC = 0.143).

#### Data S1-6

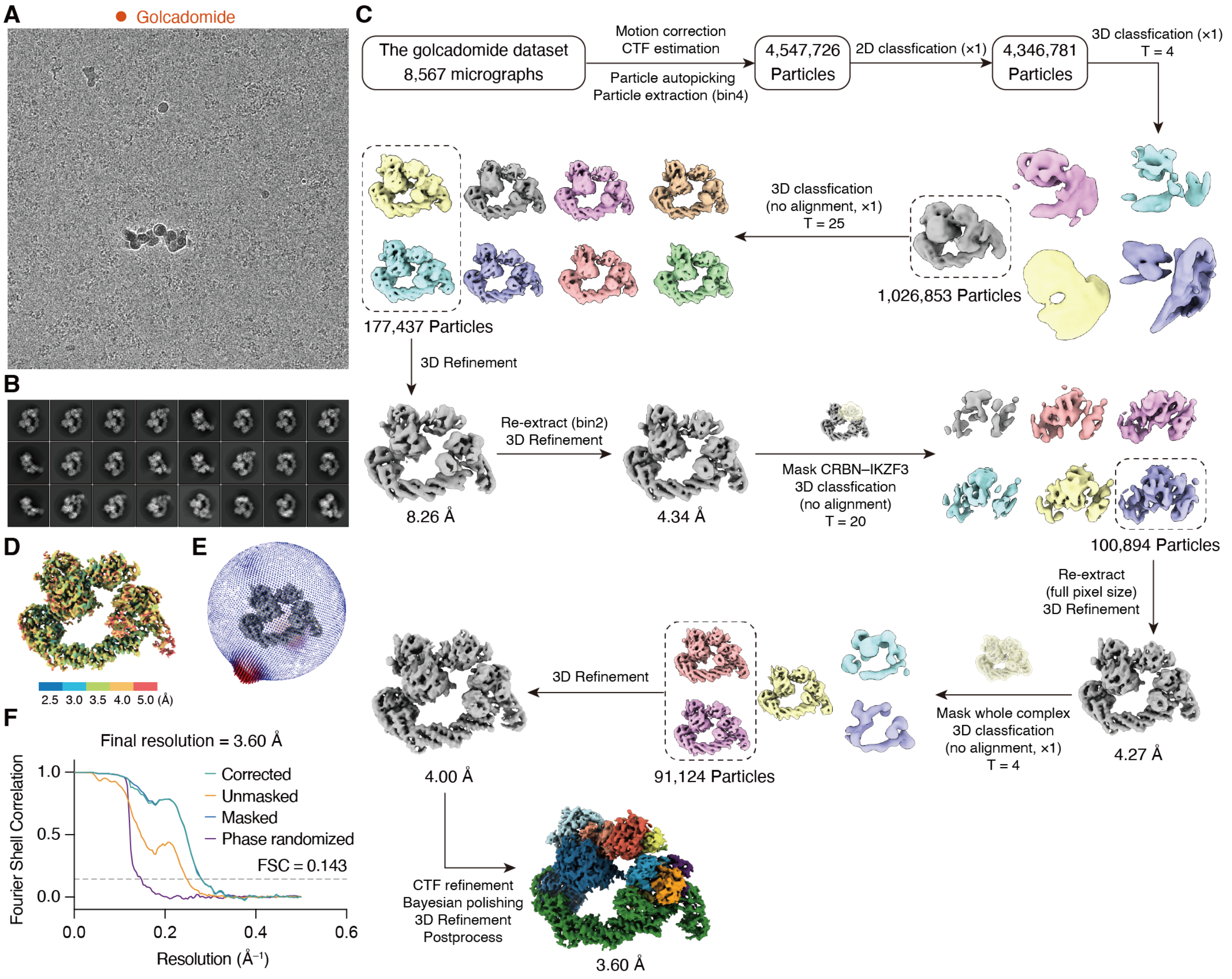


**Data S1-6. Cryo-EM data processing flow chart of the golcadomide-organized ubiquitylation assembly.** (**A**) A representative micrograph. (**B**) Representative 2D averages. (**C**) Data processing workflow. (**D**) Final cryo-EM map colored by the indicated local resolution. (**E**) Angular distributions of the final cryo-EM reconstruction. (**F**) Gold standard Fourier shell correlation (FSC) curves for the final refinement (3.60 Å resolution at FSC = 0.143).

#### Data S1-7

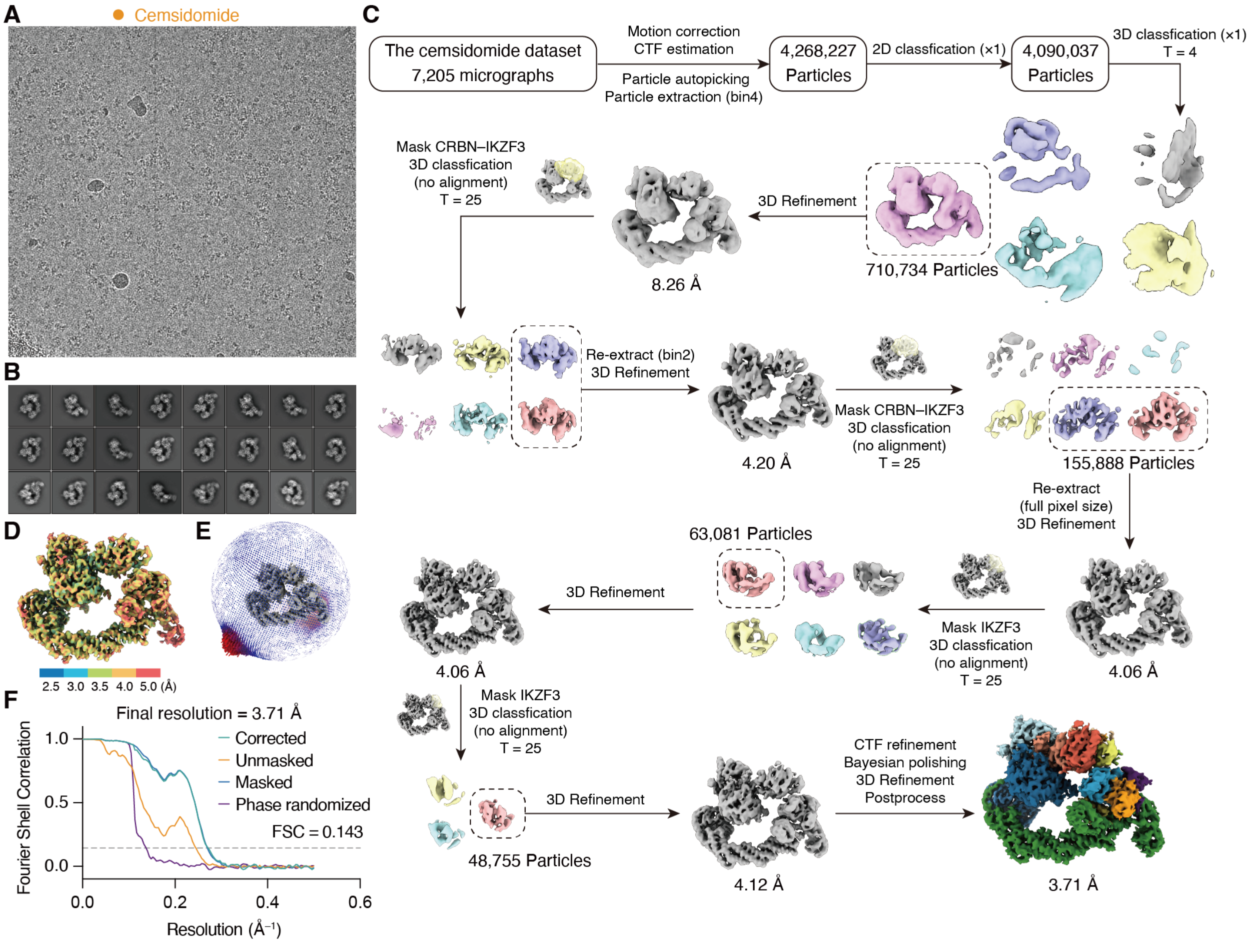


**Data S1-7. Cryo-EM data processing flow chart of the cemsidomide-organized ubiquitylation assembly.** (**A**) A representative micrograph. (**B**) Representative 2D averages. (**C**) Data processing workflow. (**D**) Final cryo-EM map colored by the indicated local resolution. (**E**) Angular distributions of the final cryo-EM reconstruction. (**F**) Gold standard Fourier shell correlation (FSC) curves for the final refinement (3.71 Å resolution at FSC = 0.143).
